# PfPHAST: *Plasmodium falciparum* Public Health Amplicon Sequencing Tool, a Streamlined Panel for Malaria Genomic Surveillance

**DOI:** 10.64898/2026.09.26.754026

**Authors:** Francis D. Semakuba, Bienvenu Nsengimaana, Monica Mbabazi, Thomas Katairo, Kathryn Murie, Nicholas Hathaway, Innocent Wiringilimaana, John Vester Muwanika, Kore Lum, Milya Davlieva, Victor Asua, Shahiid Kiyaga, Stephen Tukwasibwe, Jackie Nakasaanya, Karen B. Achom, Alisen Ayitewala, Jerry Mulondo, Eric Watyekele, Samuel L. Nsobya, Moses R. Kamya, Bosco B. Agaba, Philip J. Rosenthal, Melissa D. Conrad, Isaac Ssewanyana, Bryan Greenhouse, Andrés Aranda-Díaz, Jessica Briggs

## Abstract

Genomic tools can support malaria control policy through surveillance of *Plasmodium falciparum* populations, tracking antimalarial drug resistance, *pfhrp2/3* deletions that compromise rapid diagnostic tests, and selection at the circumsporozoite protein (PfCSP) vaccine target, as well as through molecular correction of therapeutic efficacy studies (TES). Multiplex Amplicons for Drug, Diagnostic, Diversity, and Differentiation Haplotypes using Targeted Resequencing (MAD^4^HatTeR), a comprehensive amplicon sequencing panel covering up to 276 targets, supports these applications but is tailored to research rather than routine programmatic use. We developed *P. falciparum* Public Health Amplicon Sequencing Tool (PfPHAST), a 56-target derivative of MAD^4^HatTeR spanning drug resistance loci, *pfhrp2*/*3* deletion, PfCSP genotyping, non-falciparum species identification, and 20 high-heterozygosity microhaplotype loci for TES classification. We compared PfPHAST and MAD^4^HatTeR using laboratory strain controls, including two-strain dilution series and a five-strain mixture, across parasite densities of 100 to 10,000 parasites/µL. At matched per-target depth, PfPHAST achieved a higher quality-control pass rate than MAD4HatTeR (94.4% versus 90.0%) and distributed reads more evenly across targets. The panels showed comparable recall and precision for drug resistance codons and microhaplotypes, reaching near-complete recall above 40% within-sample allele frequency (WSAF) at all densities, with reduced sensitivity for minor alleles below 10% WSAF at low parasite density in both panels. Observed and expected WSAF correlated strongly for both panels, and both resolved a five-strain polyclonal mixture, including a 5% minor strain. By concentrating sequencing capacity on targets of greatest programmatic relevance, PfPHAST offers a scalable, lower-cost alternative to comprehensive research panels without sacrificing performance on shared targets, complementing MAD^4^HatTeR for routine molecular malaria surveillance.

## Introduction

Effective malaria control depends on timely, actionable data on parasite biology, epidemiology and evolution. Genomic surveillance of *Plasmodium falciparum* can address multiple public health needs simultaneously: monitoring the emergence and spread of markers associated with antimalarial drug resistance; detecting deletions in histidine-rich protein 2 and 3 genes (*pfhrp2*/*3*) that undermine rapid diagnostic tests (RDTs); and genotyping the circumsporozoite protein (PfCSP), the target antigen of the RTS,S/AS01 and R21/Matrix-M malaria vaccines, to evaluate selective pressure and potential immune escape.^1,2^ These threats are global in scope but could have particularly profound consequences in sub-Saharan Africa, which bears the overwhelming majority of the global malaria burden.^3^ Artemisinin partial resistance has emerged in Africa and is spreading and *pfhrp2*/*3* deletions have been documented across multiple African countries.^3–5^ Meanwhile, malaria vaccines are being rolled out at scale, creating a need for surveillance to monitor their impact and effectiveness.^6^ The Africa Centres for Disease Control and Prevention (Africa CDC) has identified the monitoring of antimalarial drug resistance markers and *pfhrp2*/*3* deletions as the highest-priority use cases for continent-wide genomic surveillance;^2^ a prioritisation aligned with the World Health Organization (WHO) strategy to respond to antimalarial drug resistance in Africa.^7^ As these applications move from research settings toward routine malaria molecular surveillance (MMS), demand has intensified for genomic tools adapted to the operational realities of national malaria programmes (NMPs) and the reference laboratories that support them.

Therapeutic efficacy studies (TES) are the primary means by which antimalarial drug efficacy is monitored under programmatic conditions, and their results directly inform treatment policy.^8^ TES outcomes depend critically on the accurate classification of recurrent parasitaemia as either recrudescence or new infection, a distinction that requires genotyping of paired samples from the same patient, referred to as molecular correction. The current WHO-recommended standard relies on capillary electrophoresis of multiallelic length polymorphisms interpreted using allele match-counting.^8,9^ This approach has well-documented limitations in high-transmission settings, where polyclonal infections are common and the discriminatory power of a small number of markers is insufficient to reliably distinguish recrudescence from new infection.^10,11^ The 2021 WHO informal consultation on genotype correction methodology acknowledged these limitations and identifies amplicon deep sequencing as a medium-term target for evaluation and potential adoption as the standard approach.^9^ Microhaplotypes (two or more single nucleotide polymorphisms contained within a single amplicon) are multiallelic, and therefore contain more information per locus than biallelic SNPs. This increases the resolution with which distinct parasite clones can be distinguished within an infection and the statistical power to detect relatedness between parasites.^12,13^ A recent community-driven target product profile exercise further established that amplicon sequencing of microhaplotypes, with a sufficient number of diverse, multiallelic loci, is the recommended direction for accurate and reproducible TES genotype correction.^14^ A practical implementation of these recommendations requires a panel that combines resistance marker genotyping with a set of microhaplotype loci suitable for TES classification, within a single operationally feasible workflow.

Targeted amplicon sequencing has emerged as the preferred platform for *P. falciparum* genomic surveillance across these applications, with multiple tools now in use by laboratories across sub-Saharan Africa.^13,15,16^ More broadly, the field is converging toward a common set of public health targets regardless of sequencing platform.^17^ Among the available amplicon panels, MAD^4^HatTeR (Multiplexed Amplicons for Drug, Diagnostic, Diversity, and Differentiation Haplotypes using Targeted Resequencing) is a comprehensive Illumina-based modular panel covering as many as 276 targets, of which 243 are used in its routine configuration, including 165 high-diversity microhaplotypes, 111 antimalarial drug and diagnostic resistance targets, and PfCSP and additional vaccine targets.^13^ MAD^4^HatTeR offers broad versatility for research and surveillance applications, and has demonstrated higher sequencing depth and recall than other approaches, such as molecular inversion probe-based panels, particularly in samples with low parasite density and high polyclonality.^18^ However, the full panel requires substantial sequencing capacity while generating more data than is needed for routine surveillance and decision making. A more focused panel could retain or improve the performance characteristics of MAD^4^HatTeR (recall, precision, and microhaplotype resolution) while prioritising sequencing capacity on targets with the greatest public health utility and integrating resistance surveillance with TES classification in a single workflow.

Here, we describe the *P. falciparum* Public Health Amplicon Sequencing Tool (PfPHAST), a streamlined panel derived from MAD^4^HatTeR and tailored to the highest-priority parasite genomic surveillance needs of NMPs. By concentrating sequencing resources on drug resistance markers of programmatic relevance, *pfhrp2*/*3* deletion assessment, PfCSP genotyping, and microhaplotype loci for TES classification, PfPHAST is designed to generate data that are directly actionable by NMPs, without the overhead of a comprehensive research panel. We report the rationale and target composition of PfPHAST and characterise the performance of its microhaplotype and drug resistance targets using laboratory controls spanning a range of parasite densities and polyclonality. Assessment of *pfhrp2/3* deletions via copy number variation analysis and of performance at lower parasite densities is ongoing.

## Methods

### Panel design

PfPHAST is a derivative of MAD^4^HatTeR, designed to support routine molecular surveillance of drug and diagnostic resistance markers, and TES. The panel comprises 56 targets, against 243 in MAD^4^HatTeR, with a further four available in an optional add-on pool (described below). Targets were drawn from the parent panel, redesigned, or newly added, and organised into two primer pools (M1.1 and M2.1) to allow tiling of overlapping amplicons. Primer pools are versioned, with the suffix denoting the pool version. The assay should be run with the latest pools version, which at the time of this evaluation were M1.1 and M2.1. Updated pools will be released as targets are added or redesigned. The routine MAD^4^HatTeR configuration comprises pools D1.1 and R1.2 in the first reaction and R2.1 in the second. The PfPHAST two-pool structure requires two independent multiplexed PCR reactions (as in MAD^4^HatTeR), which are then combined into a single tube and processed through identical downstream library preparation steps (**Figure 1A**). Amplification primers were designed under contract by Paragon Genomics, Inc. using their proprietary algorithm, using the *P*. *falciparum* 3D7 reference genome, with screening against related *Plasmodium* species and the human genome to ensure specificity. Genome versions and their GeneBank or RefSeq accessions for each species are: *P*. *falciparum* Pf3D7 (version□=□2020-09-01, GCA_000002765.3), *P*. *vivax* PvP01 (version□=□2018-02-28, GCA_900093555.2), *P*. *malariae* PmUG01 (version□=□2016-09-19, GCA_900090045.1), *P*. *ovale* PocGH01 (version□=□2017-03-06, GCA_900090035.2), *P*. *knowlesi* PKNH (version□=□2015-06-18, GCA_000006355.2) and *Homo sapiens* GRCh37 (GCA_000001405.14). Full target coordinates, primer sequences, and per-target design notes are provided in **Supplementary Table 1** and **Supplementary File 1 - Note 1**.

**Figure 1.**
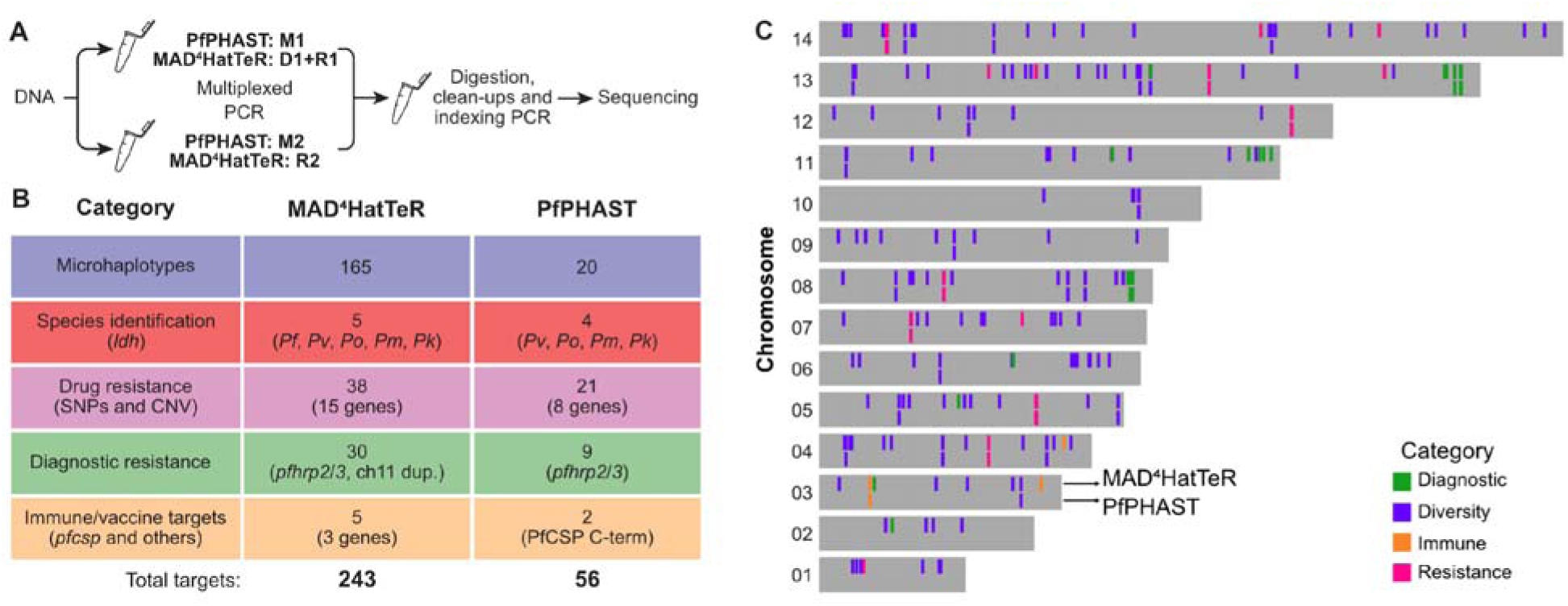
PfPHAST is a streamlined derivative of the MAD^4^HatTeR malaria amplicon sequencing panel. **A.** Simplified workflow for library preparation and sequencing. The two multiplexed PCR reactions use compatible primer pools that allow tiling of overlapping amplicons, and are combined into a single tube for all downstream steps. Pools 1 and 2 correspond to the two reactions. The pool versions used here were D1.1 and R1.2 (reaction 1) and R2.1 (reaction 2) for MAD^4^HatTeR, and M1.1 and M2.1 for PfPHAST. **B.** Target composition of the MAD^4^HatTeR and PfPHAST panels by category. Parenthetical labels indicate the genes, regions or assay types covered. Species identification targets are *ldh* in both panels: MAD^4^HatTeR includes one target per species for *P. falciparum* (*Pf*), *P. vivax* (*Pv*), *P. malariae* (*Pm*), *P. ovale* (*Po*), and *P. knowlesi* (*Pk*), while PfPHAST omits the non-informative *Pf* target and covers the remaining four species. *The optional PfPHAST M1.addon pool adds four mitochondrial *cytb* targets for the same four species. In the Diagnostic resistance category, MAD^4^HatTeR additionally targets a chromosome 11 duplication that co-occurs with *pfhrp3* deletion.^23^ **C.** Chromosomal locations of *P. falciparum* targets in the MAD^4^HatTeR (upper row) and PfPHAST (lower row) panels. Non-falciparum targets are not shown.

PfPHAST covers five target categories: antimalarial drug resistance markers; *pfhrp2*/*3* deletion assessment; the C-terminal domain of PfCSP; non-falciparum *Plasmodium* species identification; and microhaplotype loci for TES outcome classification. Selection criteria for each category are described below.

Drug resistance targets were selected to cover resistance markers with programmatic relevance in *pfkelch13* (PF3D7_1343700), dihydropteroate synthase (*pfdhps*, PF3D7_0810800), dihydrofolate reductase (*pfdhfr*, PF3D7_0417200), chloroquine resistance transporter (*pfcrt*, PF3D7_0709000), multidrug resistance protein 1 (*pfmdr1*, PF3D7_0523000), and *pfcoronin* (PF3D7_1251200) genes; and *pfmdr1* and plasmepsin II/III (PF3D7_1408000 and PF3D7_1408100) copy number variation (**Table 1**). For *pfkelch13*, targets covered all mutations classified as validated or candidate markers of artemisinin partial resistance at the time of design; the WHO compendium of markers was published subsequently.^17^ Markers included in MAD^4^HatTeR with insufficient evidence for routine programmatic genotyping were excluded (see **Supplementary File 1 - Note 1**). Five *pfkelch13* targets covering the propeller domain were redesigned with new primers to extend coverage to previously missed amino acid positions, and a new *pfmdr1* target was added to improve copy number estimation.

**Table 1:**
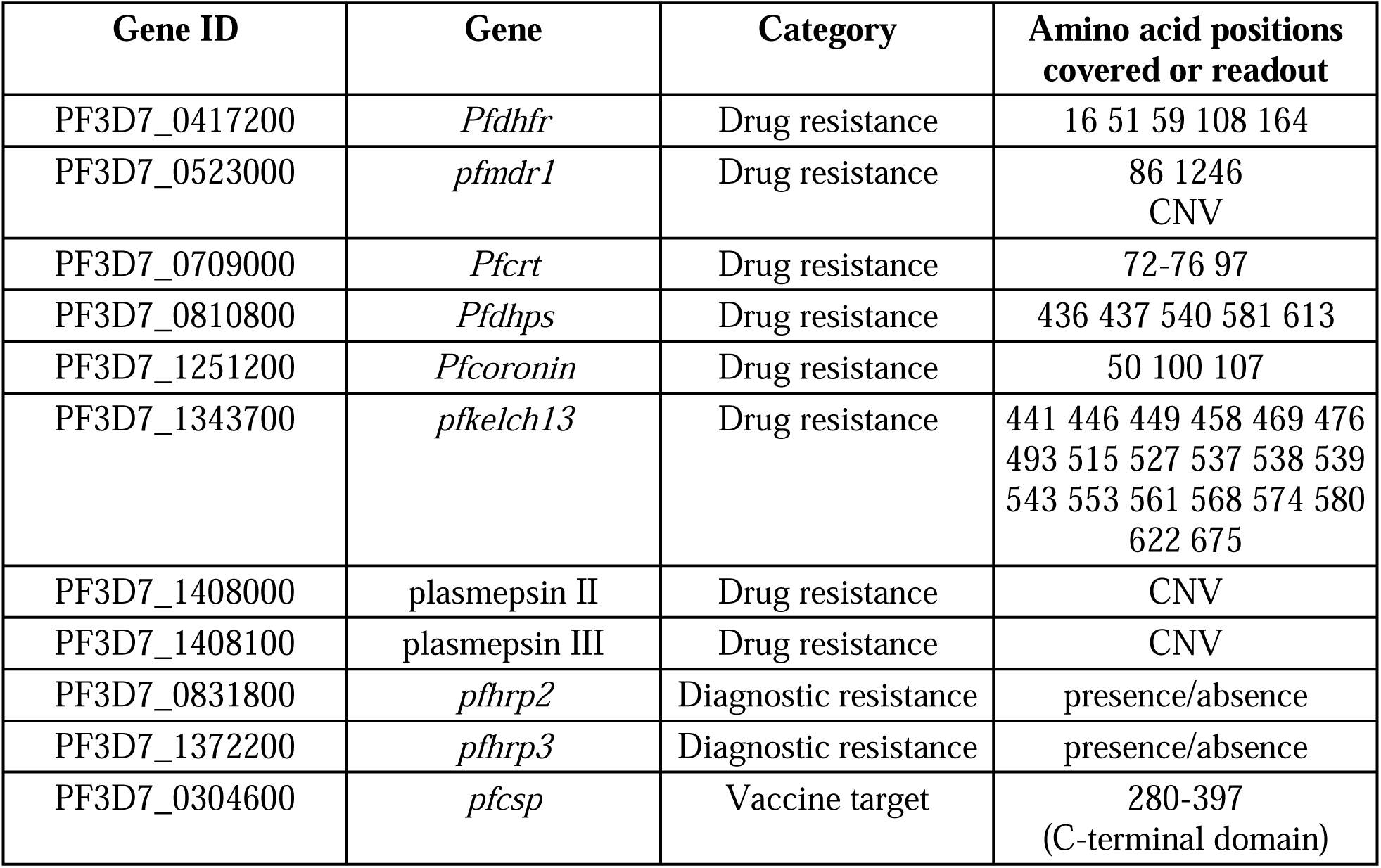
Drug resistance, diagnostic, and immune targets covered by PfPHAST. Codons listed for drug resistance genes are those classified by WHO as validated, candidate or potential markers that fall within PfPHAST amplicons.^17^ *pfmdr1*, plasmepsin II and III are assessed for copy number variation (CNV). *pfhrp2* and *pfhrp3* are assessed for gene deletion from relative read depth across targets within and flanking each gene. Full amplicon coordinates are given in **Supplementary Table 1**.

For *pfhrp2*/*3* deletion assessment, targets were retained from MAD^4^HatTeR, excluding those that consistently underperformed in the published validation data, and two new targets downstream of *pfhrp2* were added to improve deletion breakpoint resolution. Two PfCSP targets covering the Th2R, Th3R, and DV10 epitopes of the C-terminal coding domain were also retained.

Species identification targets were retained from the MAD^4^HatTeR *ldh* set, excluding the *P. falciparum* target, as *P*. *falciparum* presence is established by the rest of the panel. In addition, an optional add-on pool (M1.addon) was designed to be spiked into the first reaction alongside pool M1.1, providing increased recall for non-falciparum detection through four mitochondrial cytochrome b (*cytb*) targets, one specific to each of *P*. *vivax*, *P*. *malariae*, *P*. *ovale*, and *P*. *knowlesi*. The mitochondrial genome is present at a high copy number per parasite, yielding more templates per unit of parasite DNA than the single-copy nuclear *ldh* targets. The *P. malariae* target is predicted also to amplify *P. ovale wallikeri* (see **Supplementary File 1 - Note 1**).

Microhaplotype loci for TES outcome classification were selected from the MAD^4^HatTeR diversity module (primer pool D1.1). Twenty loci were targeted to meet the minimum specifications for TES genotype correction established by a recent community target product profile exercise, which determined through simulation that approximately 20 loci with high heterozygosity are sufficient to achieve 95% sensitivity and specificity for distinguishing recrudescence from new infection across a range of complexity of infection values.^14^ Candidate loci were restricted to those meeting performance criteria in the published MAD^4^HatTeR validation data: dropout in less than 1% of the samples (*N* = 1,103) and precision above 97%. Expected heterozygosity was estimated for each eligible locus from local haplotype reconstruction of publicly available whole-genome sequencing data as of November 2023 from Ghana, Kenya, Mali, Mozambique, and Tanzania, and loci were ranked by the minimum value across these five countries, favouring loci informative in all five populations rather than highly diverse in only some.^19,20^ Loci were then selected sequentially in order of decreasing minimum heterozygosity, provided they were at least 200 kb from any already-selected locus on the same chromosome to limit linkage between selected loci, until 20 loci were reached. Selected loci were split evenly between the two primer pools to balance the number of amplicons per reaction.

### Preparation of control samples

Controls were prepared using well-characterised *P*. *falciparum* laboratory strains: 3D7, D10, W2, D6, HB3, FCR3, V1S, and U659. D10 lacks *pfhrp2* and HB3 lacks *pfhrp3*, allowing these controls to serve as positive controls for *pfhrp2*/*3* deletion detection. Three categories of controls were included. First, monoclonal controls were prepared from 3D7, D10, and HB3. Second, two-strain dilution series controls were created by mixing pairs of strains at defined proportions. In one series, D10 was mixed at 1%, 2%, 5%, 10%, 40% and 60% with W2 making up the remainder; in the other, D6 was mixed at the same proportions with HB3 making up the remainder. These give the corresponding expected within-sample allele frequencies (WSAFs) at targets where the two strains differ. Third, a five-strain control was prepared by combining D6 (50%), V1S (20%), FCR3 (15%), W2 (10%), and U659 (5%).

All strains were maintained in continuous culture in RPMI-1640 medium as previously described.^18^ Prior to mixing, ring-stage parasites were synchronised by sorbitol lysis and quantified by Giemsa-stained thin blood smears. Strains were combined volumetrically at the target proportions, then serially diluted in whole blood from an uninfected donor to achieve parasite densities of 100, 1,000 and 10,000 parasites/µL. Aliquots of approximately 50 µL of each mixture were spotted onto Whatman 903 filter paper (Cytiva, Marlborough, MA), air-dried overnight, and stored at -20 °C until DNA extraction. These dried blood spots (DBS) were used for all subsequent processing.

### Sample processing and sequencing

DNA was extracted from DBS using the Chelex-Tween 20 method as previously described.^21^ Briefly, 6 mm discs were washed overnight in PBS (Corning 21-040-CV, Corning, NY) with 0.05% Tween 20 (Sigma Aldrich P9416, St. Louis, MO) at 4 °C, and DNA was eluted in 150 µL of 7% Chelex-100 resin (Bio-Rad, Hercules, CA) at 95 °C. The DNA-containing supernatant was transferred to barcoded cryogenic storage tubes (Micronic, Lelystad, the Netherlands), avoiding carryover of Chelex resin, and stored at -20 °C until use. Parasite density was estimated by quantitative real-time polymerase chain reaction (qPCR) targeting the var gene acidic terminal sequence (varATS) as previously described, with densities interpolated from a standard curve of 3D7 DNA serially diluted from 10,000 to 1 parasites/µL.^22^

Libraries were prepared for both MAD^4^HatTeR and PfPHAST from the same DNA extractions following the MAD^4^HatTeR protocol as previously described,^13^ using CleanPlex reagents (Paragon Genomics, Fremont, CA), with modifications described below. The complete protocol is provided as a standard operating procedure with an illustrated step-by-step guide for laboratories implementing the assay (**Supplementary File 2**).

Multiplexed PCR was performed in two reactions with compatible primer pools (MAD^4^HatTeR: reaction 1, pools D1.1 and R1.2; reaction 2, pool R2.1; PfPHAST: reaction 1, pool M1.1; reaction 2, pool M2.1) (**Figure 1A**), using 0.5 µL of each primer pool 5X solution and 15 cycles for samples with ≥100 parasites/µL, and 0.25 µL and 20 cycles for samples with <100 parasites/µL. The two reactions were then combined into a single tube and processed together through all subsequent steps. CleanMag magnetic bead (Paragon Genomics) clean-ups, digestion of non-specific products, and indexing PCR were performed following the CleanPlex protocol, with 80% ethanol for all CleanMag magnetic bead washes. Indexing used custom 12 bp TruSeq indexes (Integrated DNA Technologies, Coralville, IA) and CleanPlex indexes (10 bp, Paragon Genomics, Fremont, CA), which are compatible in a single run. The full set of twelve custom and four CleanPlex 96-index plates provides 1,536 combinations, of which a subset was used here.

Libraries were prepared manually with 8-channel pipettes in 96-well plate batches, each including 1 to 2 positive (3D7 DNA at 1,000 parasites/μL) and 2 to 6 negative controls (nuclease-free water). Plates were shared with field samples from Northern Uganda, reflecting routine laboratory practice in which controls are processed within batches of clinical samples rather than in isolation. Library quality and fragment size distribution were assessed by capillary electrophoresis (Agilent Bioanalyzer or TapeStation, Agilent Technologies, Santa Clara, CA) for all positive and negative controls and for two randomly selected samples per plate. Absence of a peak at approximately 400 bp in the negative controls was confirmed before proceeding to ensure there was no contamination during library preparation.

After indexing PCR, libraries were pooled at volumes inversely proportional to parasite density to balance read counts per sample (15 µL for 100 to <1,000, 10 µL for 1,000 to <10,000, 6 µL for 10,000 to <100,000, and 3 µL for ≥100,000 parasites/µL). Pools were cleaned at a 1X CleanMag bead-to-library ratio, eluted in 43 µL of Tris-EDTA buffer, assessed by capillary electrophoresis and quantified with a Qubit Flex (Thermo Fisher Scientific, Waltham, MA).

The number and arrangement of replicates differed between panels. Each MAD^4^HatTeR control was prepared in duplicate on one plate and re-prepared in duplicate from the same DNA extract on a second plate, sequenced in a separate run, giving four independent library preparations per control. PfPHAST controls were prepared in duplicate, either as two independent preparations sequenced in separate runs, or as two preparations within a single plate sequenced on the same run.

Libraries were sequenced on an Illumina MiSeq or NextSeq 2000 (Illumina, San Diego, CA) using a 600-cycle v3 kit and a 300-cycle P1 kit, respectively. Both used 151 bp paired-end reads, with PhiX spiked in at 5% for the MiSeq and 2% for the NextSeq 2000. MAD^4^HatTeR libraries were sequenced on the MiSeq only; PfPHAST libraries were sequenced on both platforms.

### Data analysis

Sequencing data were analysed with the MAD^4^HatTeR bioinformatics pipeline v1.0.0,^24^ which we adapted to work with any panel configuration. The pipeline returns allele-level outputs comprising sample identifier, locus, allele sequence (as a pseudoCIGAR string) and read count, together with amino acid calls and counts at predefined drug resistance markers. These were imported into R version 4.6.0 (R Foundation for Statistical Computing, Vienna, Austria) together with per-sample metadata on parasite density and replicate. All downstream analyses were performed in R. Both panels were processed with the same pipeline and parameters. To account for the lower per-target depth expected for MAD^4^HatTeR, which comprises 243 targets against 56 for PfPHAST, reads were pooled across pairs of MAD^4^HatTeR replicate libraries, giving two observations per control for each panel and comparable per-target depth.

Two categories of targets were excluded before analysis, applied identically to both panels. The *ldh* targets used for species identification were uninformative here, as all controls consisted of *P. falciparum* strains. Because longer amplicons amplify less efficiently and consistently yield lower read depth, targets of more than 275 bp were also excluded. 50 of the 56 PfPHAST targets and 229 of the 243 MAD^4^HatTeR targets were retained. Samples in which fewer than 75% of the retained targets reached at least 100 reads failed quality control (QC) and were excluded. The proportion of samples passing QC was calculated for each parasite density stratum. In the samples passing QC, read depth was summarised per target as the median and interquartile range across samples, and per sample as the mean across retained targets, stratified by parasite density. Alleles with fewer than 10 reads or a WSAF below 0.75% were excluded, and WSAF was recalculated across the remaining alleles at each target.

Detection performance was assessed as recall (also termed sensitivity, the proportion of expected alleles that were detected), and precision (the proportion of detected alleles that were expected). Expected alleles were taken from sequencing of monoclonal controls of the eight strains, reported previously: at each target, the expected alleles of a control were the genotypes of its constituent strains, each with an expected WSAF defined by its mixing proportion. Alleles were compared at two levels: microhaplotypes and the amino acid encoded at each drug resistance codon. Of the 75 codons in the WHO compendium falling within the genes covered by either panel, 57 are targeted by both. Amino acid calls were compared at the subset of these that differ between the constituent strains of at least one control and are therefore informative for detection, so that both panels were evaluated on the same codons. Microhaplotypes were compared at all informative TES-classification microhaplotypes and PfCSP targets of each panel, which differ between panels. Targets and codons lacking a true genotype, or at which all strains carried the same allele, carried no information and were excluded. Recall and precision were calculated from counts of true positives (TP, alleles both observed and expected), false positives (FP, observed but not expected) and false negatives (FN, expected but not observed), as TP/(TP+FN) and TP/(TP+FP), and summarised by panel and parasite density. Recall was additionally summarised by the WSAF. The accuracy of WSAF estimation was assessed as the squared Pearson correlation (r^2^) between observed and expected WSAF within each panel and parasite density, with undetected expected alleles assigned an observed WSAF of zero and targets with fewer than 100 reads in a given library excluded.

To test whether recall differences in diversity and immune microhaplotypes reflected panel performance rather than marker content, we restricted both panels to shared microhaplotype targets. This yielded 22 diversity and immune loci with an identical set of 465 expected alleles per panel. At each parasite density and expected WSAF level, we computed the recall difference (PfPHAST − MAD^4^HatTeR), with 95% confidence intervals from 2,000 bootstrap replicates resampling libraries with replacement within density strata.

## Results

### PfPHAST panel composition

PfPHAST comprises 56 targets organised into two primer pools (M1.1, *n* = 41; M2.1, *n* = 15), compared with 243 in the routine MAD^4^HatTeR configuration (**Figure 1B**). The panel comprises (i) 20 microhaplotype loci with high expected heterozygosity (minimum heterozygosity > 0.71 across five reference populations, **Supplementary Table 2**), for genotyping in TES; (ii) four *ldh* targets for identification of non-falciparum *Plasmodium* species (*P. vivax*, *P. malariae*, *P. ovale*, and *P. knowlesi*), with four mitochondrial *cytb* targets for the same species available in the optional M1.addon pool; (iii) 21 drug resistance targets across eight genes (*pfkelch13*, 5; *pfmdr1*, 4; *pfdhps*, 3; *pfdhfr*, 2; *pfcoronin*, 2; *pfcrt*, 1; plasmepsin II and plasmepsin III, 4 for copy number); (iv) 9 targets for *pfhrp2*/*3* deletion detection (4 in and around *pfhrp2*, 5 in and around *pfhrp3*); and (v) 2 targets in the PfCSP C-terminal domain, for monitoring selective pressure and potential immune escape at the vaccine target. Targets are distributed across 12 of the 14 chromosomes (**Figure 1C**). As in MAD^4^HatTeR, the two primary pools are amplified in separate reactions and combined for all subsequent steps (**Figure 1A**).

### Sequencing depth and quality control

Fifteen mixed controls (14 two-strain controls and one five-strain) were assessed in duplicate with each panel across three parasite densities (100, 1,000 and 10,000 parasites/μL); this resulted in 30 control samples per panel per parasite density. MAD^4^HatTeR libraries received more total reads per sample than PfPHAST (**Supplementary File 1 - Figure 1**), but with 243 targets against 56, per-target depth remained lower. To compare the panels at similar per-target depth rather than at similar sequencing effort, reads from pairs of MAD^4^HatTeR replicate libraries were pooled, resulting in a median per-target depth of 708 reads compared with 739 for PfPHAST. All subsequent comparisons use these depth-matched data.

Overall, 94.4% (85/90) of PfPHAST samples and 90.0% (81/90) of MAD^4^HatTeR samples across all parasite densities passed the QC criteria described in the **Methods** and were retained for downstream analysis. Reads were distributed more evenly across targets in PfPHAST than in MAD^4^HatTeR (interquartile range width of 231 versus 549 reads; **Figure 2**). Median depth exceeded 100 reads for 98.1% of PfPHAST targets and 97.0% of MAD^4^HatTeR targets, with seven MAD^4^HatTeR targets consistently below this depth.

**Figure 2.**
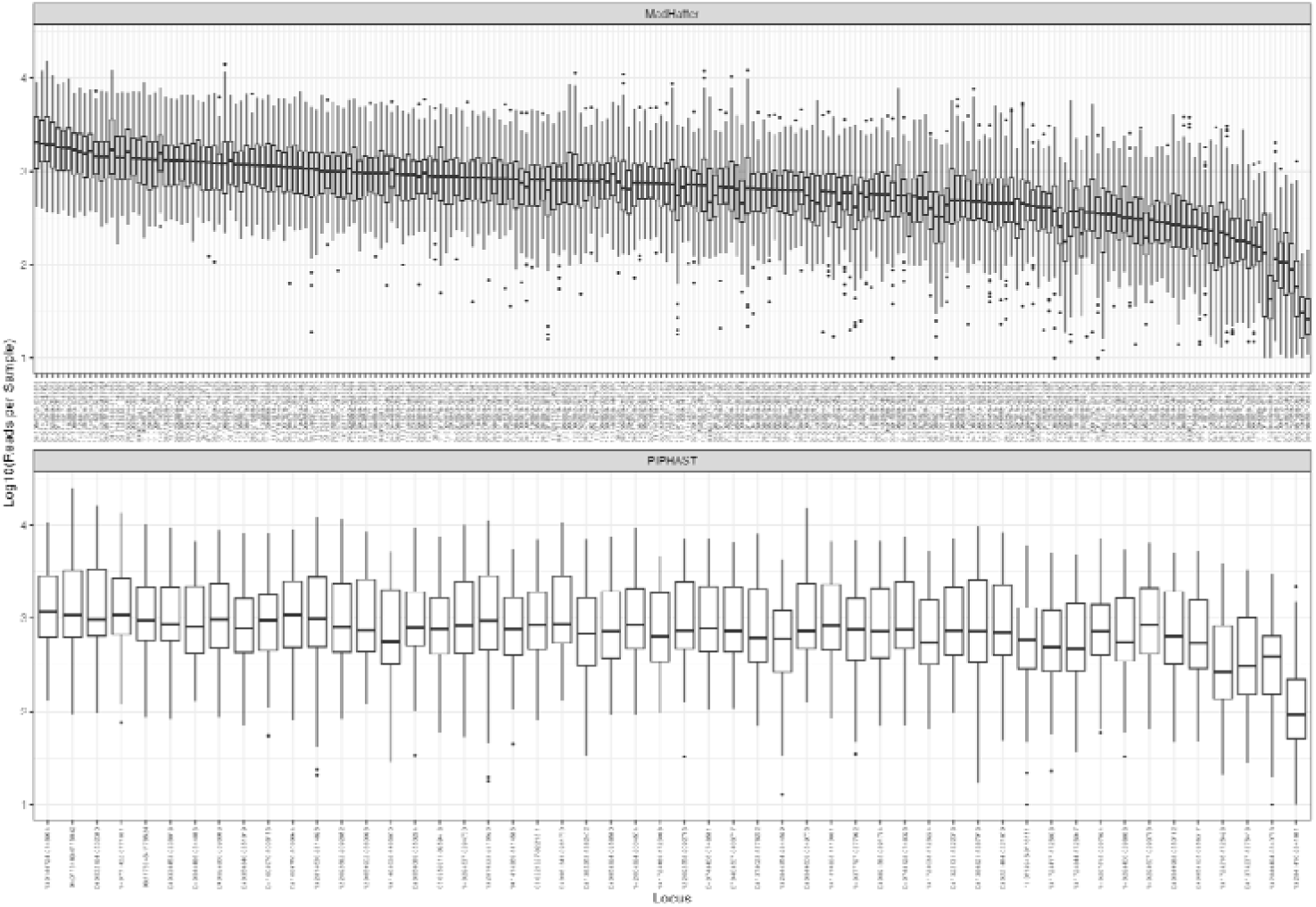
**Per-target sequencing depth for MAD**^4^**HatTeR and PfPHAST.** Boxes show the median and interquartile range across samples passing quality control from all parasite densities, with whiskers extending to 1.5× the interquartile range; targets are ordered by descending median depth independently for each panel. Reads were pooled across pairs of MAD^4^HatTeR replicate libraries to match depth between panels, as described in the **Methods**.

### PfPHAST and MAD^4^HatTeR detect drug resistance codons with comparable recall, quantification and precision

Both panels target 57 of the 75 WHO-compendium codons in the genes they cover. 10 of these differ between the constituent strains of at least one two-strain control and are therefore informative for detection at varying WSAF. Performance was compared between the two panels using the same 10 codons. Mean recall increased with expected WSAF for both panels, reaching 100% at WSAF of 40% or above at all three densities (**Figure 3A**). Recall was lowest at 100 parasites/µL for alleles below 10% WSAF, and improved at 1,000 parasites/µL for both panels; a further improvement at 10,000 parasites/µL was seen for PfPHAST but not MAD^4^HatTeR. Observed and expected WSAF were strongly correlated for both panels, with correlation increasing with parasite density for PfPHAST (**Figure 3B, C**).

**Figure 3.**
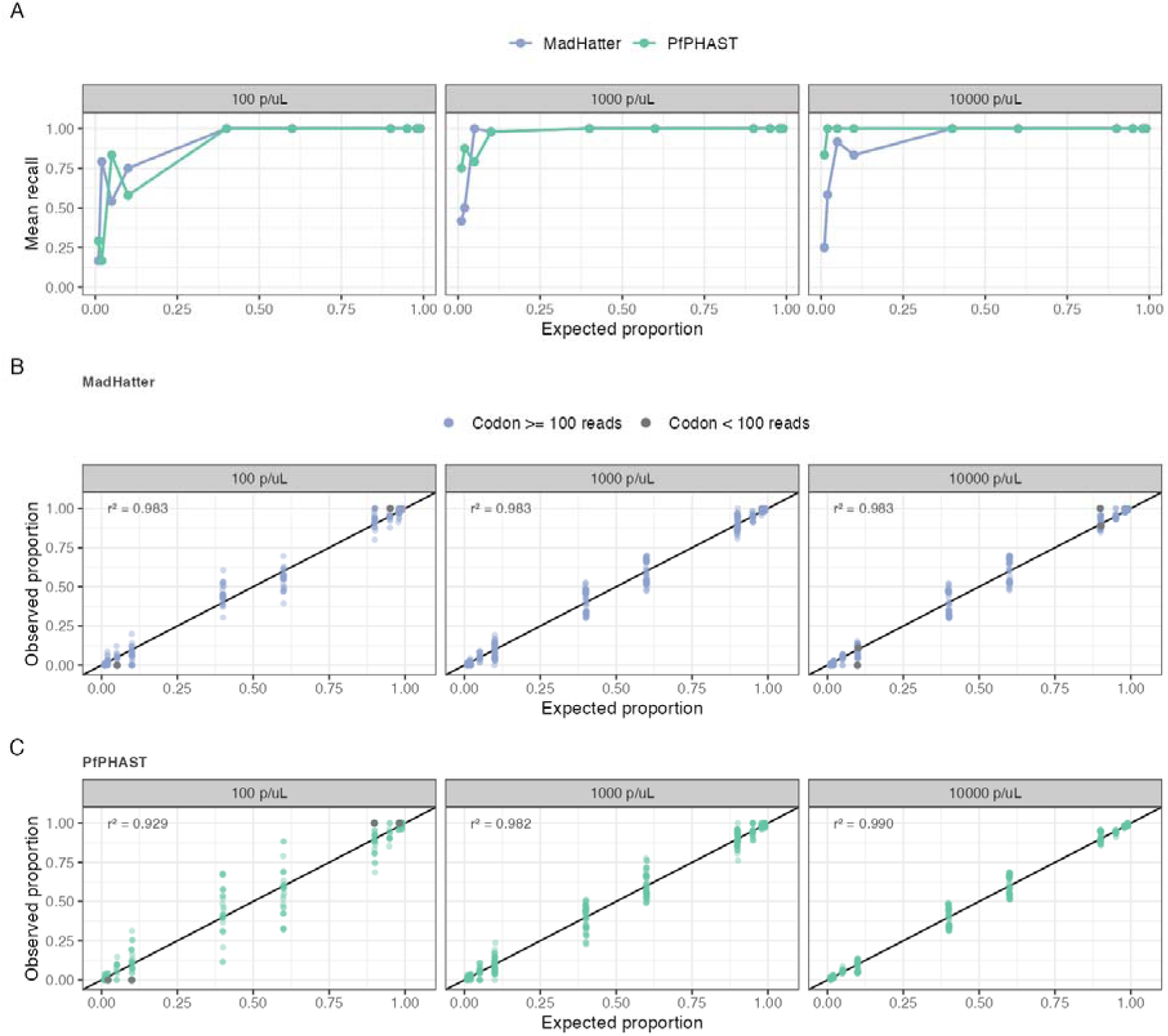
Drug resistance codon detection by MAD^4^HatTeR and PfPHAST. **(A)** Mean recall for amino acid alleles at drug resistance marker codons, averaged across expected alleles. **(B, C)** Observed versus expected within-sample allele frequency for MAD^4^HatTeR (B) and PfPHAST (C), with the black line showing y = x. Grey points indicate codons with fewer than 100 reads, which were excluded from the r^2^ calculation.

False-positive codon calls were extremely rare in both panels. Across the 45 codons where the truth was known for all strains, evaluated in all control types (two-strain dilution series, the five-strain control, and single-strain controls), PfPHAST produced one false positive in 4,145 codon calls. The call was PfKelch13 C469Y, a validated artemisinin partial resistance marker that circulates at appreciable frequency in East Africa,^21,25,26^ detected at 3.71% WSAF in a D10 single-strain control at 100 parasites/µL. This library shared a plate with field samples, 27% of which carried PfKelch13 C469Y. MAD^4^HatTeR produced none among 4,383 (**Figure 4**).

**Figure 4.**
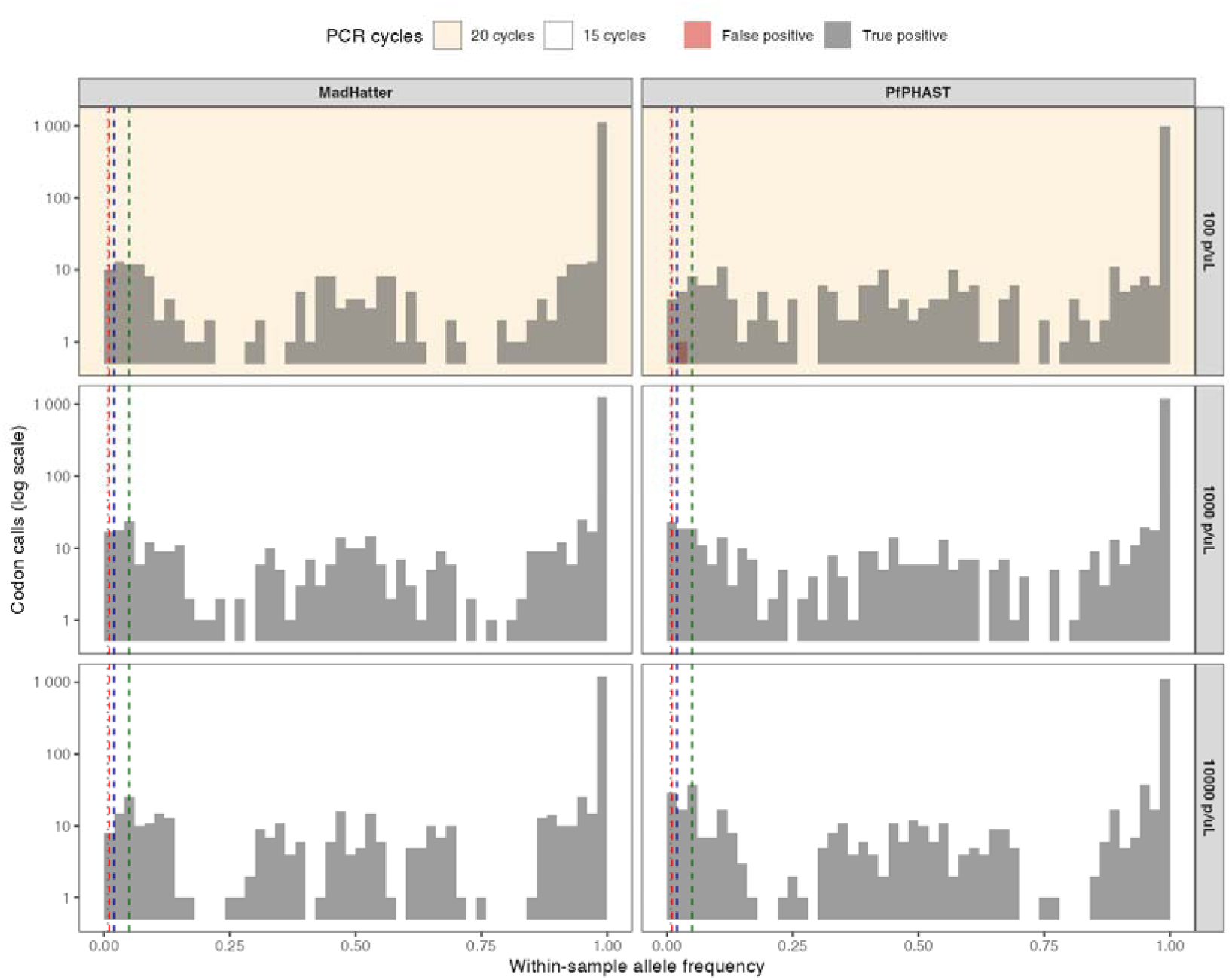
Distribution of true- and false-positive drug-resistance codon calls by within-sample allele frequency (WSAF). Data span all 45 WHO-compendium codons with control truth data, across all control types (two-strain dilution-series, the five-strain control, and the D10 and HB3 single-strain controls), with each panel scored only on the codons it targets. Calls required at least 10 reads and 0.75% WSAF. Vertical lines mark the applied frequency floor (dotted, 0.75%) and alternative thresholds at 1%, 2% and 5% (dashed).

### PfPHAST and MAD^4^HatTeR recover microhaplotypes and resolve polyclonal infections comparably

Both panels reliably recovered expected microhaplotypes in the two-strain controls, with mean recall increasing with parasite density and expected WSAF (**Figure 5A**). Recovery of minor microhaplotypes was poorest at 100 parasites/µL, where both panels detected fewer than half of those expected at 2% WSAF or below, and improved at 1,000 and 10,000 parasites/µL. Above 40% expected WSAF, both panels recovered ≥99% of expected microhaplotypes at every density.

**Figure 5.**
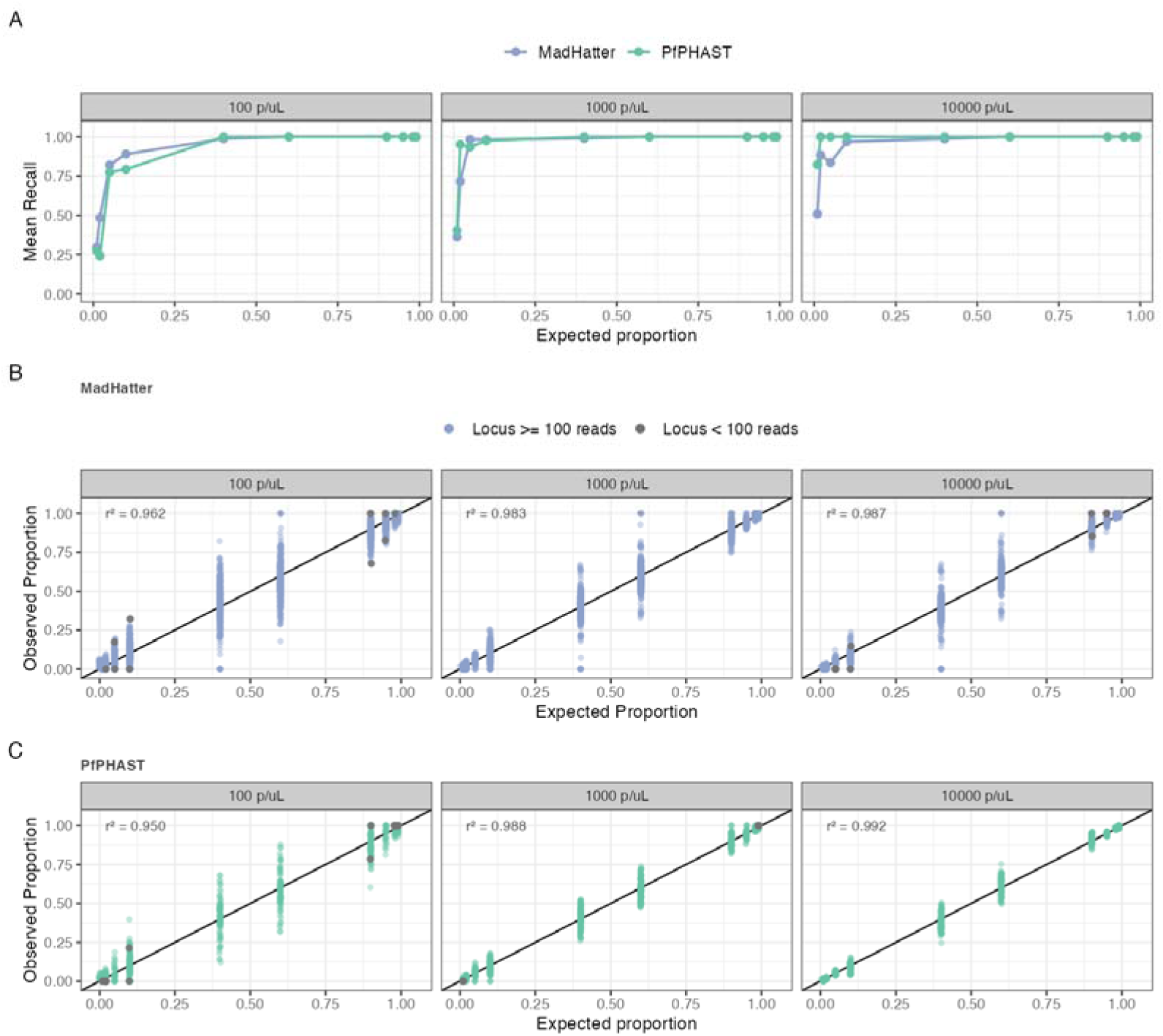
**Microhaplotype detection by MAD**^4^**HatTeR and PfPHAST. (A)** Mean recall for microhaplotype alleles, evaluated on the diversity and PfCSP targets for MAD^4^HatTeR and PfPHAST, averaged across expected alleles. **(B–C)** Observed versus expected microhaplotype within-sample allele frequency for **(B)** MAD^4^HatTeR and **(C)** PfPHAST, with the black line showing y = x. Grey points indicate targets with fewer than 100 reads, which were excluded from the r^2^ calculation.

Evaluated across all informative microhaplotype and PfCSP targets contained in each panel (122 for MAD^4^HatTeR, 22 for PfPHAST), the two performed comparably, with differences confined to the lowest expected WSAF: at 100 parasites/µL, MAD^4^HatTeR recovered more at 10% WSAF or below, while at 1,000 and 10,000 parasites/µL, PfPHAST recovered more at 1 to 2% WSAF. Restricting both panels to the 22 targets they share gave the same pattern, though at 100 parasites/µL no per-WSAF difference was significant; at 10,000 parasites/µL the differences at 1% and 2% remained significant, in favour of PfPHAST (**Supplementary Table 3**). Observed and expected WSAF were strongly correlated for both panels (**Figure 5B, C**), with correlation increasing with parasite density. False-positive microhaplotype calls were more frequent than FP drug resistance marker codon calls, and concentrated at low WSAF and 100 parasites/µL (**Supplementary File 1 - Figure 2**).

Both panels detected all five strains of the five-strain control (D6 50%, V1S 20%, FCR3 15%, W2 10%, U659 5%), including the 5% minor strain (**Figure 6**). Targets were restricted to those carrying at least four distinct alleles across the five strains, resulting in 8 targets shared by MAD^4^HatTeR and PfPHAST, and 2 additional targets unique to MAD^4^HatTeR. At most of these, two or more strains share an allele and cannot be distinguished; only two MAD^4^HatTeR targets and one PfPHAST target vary at all five. Where the 5% U659 component was resolvable it was detected by both panels, and observed proportions approximated expected proportions at 1,000 and 10,000 parasites/µL. At 100 parasites/µL both panels departed further from expected composition than at higher densities, most markedly for MAD^4^HatTeR, where no replicate passed QC and pooled depth reached only 52 to 380 reads per target against 1,007 to 1,657 for PfPHAST.

**Figure 6.**
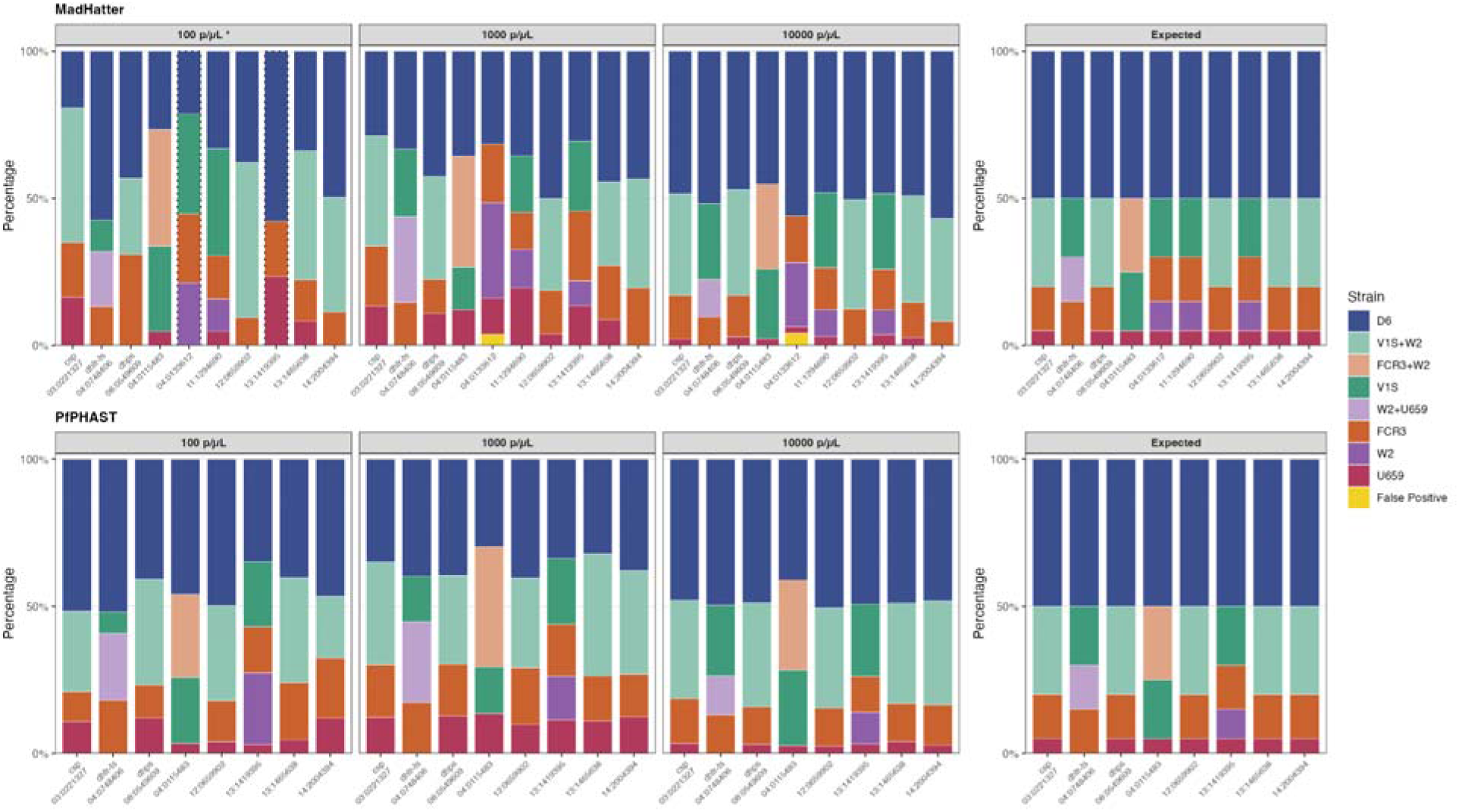
Resolution of a five-strain polyclonal control by MAD^4^HatTeR and PfPHAST. Stacked bars show the observed allele composition of the five-strain control (D6 50%, V1S 20%, FCR3 15%, W2 10%, U659 5%) by target, alongside the expected composition. Each band is one distinct allele, ordered by expected frequency with the largest at the top; reads are pooled across both replicates within each density. A band labelled with two strains is a target at which those strains share an allele, so reads cannot be attributed between them; yellow denotes alleles not expected for this control. Targets are restricted to those carrying at least four distinct alleles across the five strains. The asterisk indicates that no MAD^4^HatTeR replicate passed quality control at 100 parasites/µL, and dotted bar outlines indicate targets with fewer than 100 pooled reads.

## Discussion

PfPHAST is a purpose-built targeted amplicon sequencing panel derived from the larger MAD^4^HatTeR panel to support routine MMS by NMPs. By focusing on 56 targets selected for priority surveillance applications, PfPHAST requires roughly a quarter of the sequencing reads needed by MAD^4^HatTeR to achieve equivalent per-target depth. At 500 reads per target, a run yielding approximately 25M reads (MiSeq v3 or MiSeq i100 25M kits) accommodates approximately 884 samples, against 204 for MAD4HatTeR, allowing greater multiplexing and lower per-sample sequencing costs. The panel captures markers relevant to the highest-priority MMS use cases: monitoring antimalarial drug and diagnostic resistance, identifying non-falciparum species, tracking potential vaccine-driven selection within *pfcsp*, and classifying recurrent infections in TES.^2,9^ Here, we show robust amplification across the panel and performance comparable to MAD^4^HatTeR for shared drug resistance and microhaplotype targets, supporting the use of a more focused panel for these surveillance applications. Assessment of *pfmdr1* and plasmepsin II/III copy number and of *pfhrp2*/*3* deletion is ongoing. Because PfPHAST produces the same per-target read-depth data as MAD^4^HatTeR, the generalised additive model approach developed for that panel to estimate copy number is directly applicable.^13^

PfPHAST maintained the high performance of MAD^4^HatTeR while providing more uniform coverage across its target set. Quality control was passed by 94.4% of PfPHAST samples, compared with 90.0% for MAD^4^HatTeR at matched per-target depth, and reads were distributed more evenly across targets. Because the densities tested span the range expected for RDT-positive clinical samples, this pass rate is informative for the sample types NMPs will encounter in routine surveillance.^27,28^ Consistent amplification and sequencing depth are important not only for data quality but also for minimising repeat processing, sequencing costs, and turnaround time, all of which constrain surveillance programmes processing large numbers of samples with limited sequencing capacity.

Both panels recovered drug resistance codons and diversity microhaplotypes with complete or near-complete recall at 40% WSAF and above at every parasite density tested. They also recovered minor alleles at lower frequencies, with recall declining below 10% WSAF and most markedly at 100 parasites/µL. Observed and expected allele frequencies agreed closely for both panels. Reliable detection and quantification of resistance-associated variants underpin monitoring of changes in the prevalence of established and emerging resistance markers, and accurate recovery of diversity microhaplotypes and their WSAFs underpins TES, where these data distinguish recrudescence from new infection, a process referred to as molecular correction.^8,29^ Both panels also resolved a five-strain polyclonal control, including a strain present at 5%, indicating that minority clones can be recovered from complex infections when parasite density is adequate. Reduced recovery of minor alleles may be particularly consequential in TES, where recurrent infections often occur at low parasite density and failure to detect minority clones could affect recurrent infection classification. No available genotyping assay recovers every minor allele, however, and classification methods should therefore account for incomplete observation of the alleles present. The very low false-positive rate (1/8,528 codon calls) indicates high specificity, which is critical for resistance surveillance, where spurious detection of rare or emerging variants could prompt unnecessary concern or investigation. The single false positive occurred in a library prepared on a plate where field samples carrying PfKelch13 C469Y were common, and most plausibly reflects well-to-well contamination rather than amplification or sequencing error. Together, these findings support PfPHAST as a reliable method for routine MMS, provided parasite density and the limits of minority-allele detection are accounted for when interpreting the data.

PfPHAST complements rather than replaces MAD^4^HatTeR, providing a more focused option for routine MMS applications. MAD^4^HatTeR’s broader target set remains advantageous for research questions requiring higher-resolution genetic characterisation; for example, its larger set of microhaplotypes provides greater resolution for analyses of parasite relatedness and transmission.^13^ However, the additional loci come at the cost of greater sequencing capacity and computational and storage demands. PfPHAST addresses these constraints by concentrating sequencing capacity on a smaller set of targets prioritised for routine MMS. The trade-off is deliberate: data generated with PfPHAST will support the surveillance applications it targets, while questions requiring greater genomic resolution are better served by the broader panel. Selecting between the panels therefore requires users to define their surveillance or research objectives in advance and to determine the level of genomic resolution needed, balancing information content against sequencing and analytical requirements.

Implementation of MMS at scale depends on cost and on reliable reagent supply. Reagents for library preparation of both panels are available through Paragon Genomics, offering a common library preparation chemistry and a single supply source across the panel family. The complete protocol is provided as a standard operating procedure with an illustrated step-by-step guide for laboratories adopting the assay (**Supplementary File 2**). Costed at the Central Public Health Laboratories, Kampala, Uganda, at the time of writing and under global health pricing, the sample-to-data cost of PfPHAST, including DNA extraction, library preparation, and sequencing, is approximately US$16 per sample when 1,536 samples are multiplexed on a single NextSeq 2000 P1 run, a level of multiplexing achievable with a combination of custom and commercially available unique dual indexes. The equivalent cost for MAD^4^HatTeR is approximately US$19 per sample, reflecting the 4-fold higher per-sample read requirement of the larger panel. Because samples failing quality control incur the same cost as those that succeed, and PfPHAST had the higher pass rate, its advantage per reportable sample is larger than the difference per sample processed suggests.

PfPHAST and MAD^4^HatTeR are in use in at least 24 laboratories worldwide, including national reference laboratories in low- and middle-income countries, where the resulting data support malaria case management and inform surveillance and public health responses to the emergence and spread of drug and diagnostic resistance.^21,30–37^ PfPHAST has been adopted for the Africa CDC regional MMS initiative, through which 10 national laboratories have implemented the workflow.^38^ Laboratory protocols and bioinformatics documentation are available online,^39,40^ including an AI assistant that answers questions about the analysis workflow, lowering the barrier for laboratories without dedicated bioinformatics support. The primary bioinformatics pipeline and standardised QC code can be run locally or through Terra, supporting implementation across laboratories with different computational resources and expertise.^41^ Additional tools are available for downstream analysis, including copy number variation analysis.^42^ Ongoing work aims to simplify the laboratory workflow, reduce per-sample cost, and improve the bioinformatics pipeline, including detection of cross-contamination. Adaptation of the PfPHAST targets to other sequencing chemistries is also under way, which would extend their use to programmes without access to Illumina platforms.

As MMS priorities evolve, PfPHAST and other targeted sequencing panels will require periodic updates to incorporate newly relevant drug resistance markers and emerging diagnostic targets. The modular design of PfPHAST facilitates the rapid incorporation or redesign of targets as they become relevant, while the panel-agnostic bioinformatics pipeline allows new configurations to be accommodated without modification of the underlying code. The construction of PfPHAST itself illustrates how accumulated performance data can inform these updates: targets that underperformed in the MAD4HatTeR validation data were excluded where they were not required for a priority use case, candidate microhaplotype loci were filtered on observed dropout and precision, and targets were redesigned or added where coverage was incomplete (**Supplementary File 1 - Note 1**). Updated PfPHAST primer pools (M1.2 and M2.2) are already in development, covering markers included in the WHO compendium, which was published after the panel was designed, and newly identified candidate resistance markers.^17,43,44^ Continued monitoring of data generated by both panels will similarly help identify systematic target dropout and guide future refinement. New targets will nevertheless need to be carefully prioritised to preserve the panel’s focused and lightweight design.

Several aspects of PfPHAST performance require further evaluation. Samples below 100 parasites/µL were not analysed here, and performance in this range remains to be established. The increased number of false-positive microhaplotype calls observed at 100 parasites/µL, where libraries were amplified for 20 rather than 15 cycles, indicates that amplification conditions, bioinformatic processing, or both will require optimization for low-density samples. Copy number variation, including detection of *pfhrp2*/*3* deletions and of *pfmdr1* and plasmepsin II/III copy number, was also not evaluated here. The optional M1.addon pool for non-falciparum *Plasmodium* species identification was likewise not assessed. Replicates differed between panels, with MAD4HatTeR reads pooled across replicate libraries to match per-target depth, so the two panels were not replicated identically. Expected heterozygosity for the microhaplotype loci was estimated from African parasite populations, so their informativeness in other geographies remains to be characterised, and because the panel targets the minimum number of loci specified for TES classification,^14^ dropout of individual loci may affect classification performance. Finally, all controls were cultured laboratory strains spotted on filter paper; evaluation using real-world surveillance samples will be important to establish performance across the broader range of parasite densities, infection complexities, and sample collection and storage conditions encountered in routine use, and such evaluation is ongoing. Together, these evaluations will better define the performance and utility of PfPHAST for routine MMS.

Overall, PfPHAST provides a focused and scalable complement to MAD^4^HatTeR, prioritising *P. falciparum* genetic targets with direct relevance to public health decision making, including antimalarial drug and diagnostic resistance, the PfCSP vaccine target, and microhaplotype loci for recurrent infection classification in TES. By coupling the assay with accessible analytical tools, and by allowing new markers to be added as parasite populations and surveillance priorities evolve, PfPHAST offers a sample-to-data framework for translating parasite genomic data into routine malaria surveillance. Together, these features position PfPHAST to expand the scale and accessibility of MMS and strengthen the integration of parasite genomics into public health practice.

## Acknowledgements

We thank members of the EPPIcenter at UCSF, the Infectious Disease Research Collaboration, and the ISGlobal Malaria Physiopathology and Genomics groups for valuable discussions.

## Contributions

FDS, BN, TK, KM, NH, PJR, MDC, BG, AAD and JB conceived and designed the study. FDS, BN, TK, IW, JVM, JN, KBA, KL and MD performed the experiments. FDS, BN, KM, SK, AAD and JB analysed the data. FDS, BN, TK, MM, KM, VA, EW, BG, AAD and JB contributed to the interpretation of the results. AA, SLN, MRK, BBA, IS, ST, PJR, BG, AAD and JB supervised the study. FDS, BN, TK, MM, KM, AAD and JB drafted the manuscript. All authors reviewed the manuscript.

