## Supplementary File 1 for "PfPHAST: *Plasmodium falciparum* Public Health Amplicon Sequencing Tool, a Streamlined Panel for Malaria Genomic Surveillance"

### Supplementary materials

**Supplementary File 1 (this document)**

- **Supplementary Figure 1**. Total sequencing reads per sample for MAD$^4$HatTeR and PfPHAST.
- **Supplementary Figure 2.** Distribution of true- and false-positive microhaplotype allele calls by within-sample allele frequency.
- **Supplementary Note 1.** Per-target design notes and marker exclusions.

**Supplementary Tables** (provided as a separate file Supplementary_Tables.xlsx)

- **Supplementary Table 1.** Target coordinates and primer sequences for the PfPHAST panel.
- **Supplementary Table 2.** Expected heterozygosity, dropout and precision of the 20 TES classification microhaplotype loci.
- **Supplementary Table 3.** Differences in mean recall between panels by expected WSAF and parasite density.

**Supplementary Protocols** (provided separately)

- **Supplementary File 2.** Illustrated standard operating procedure for laboratories implementing the assay.

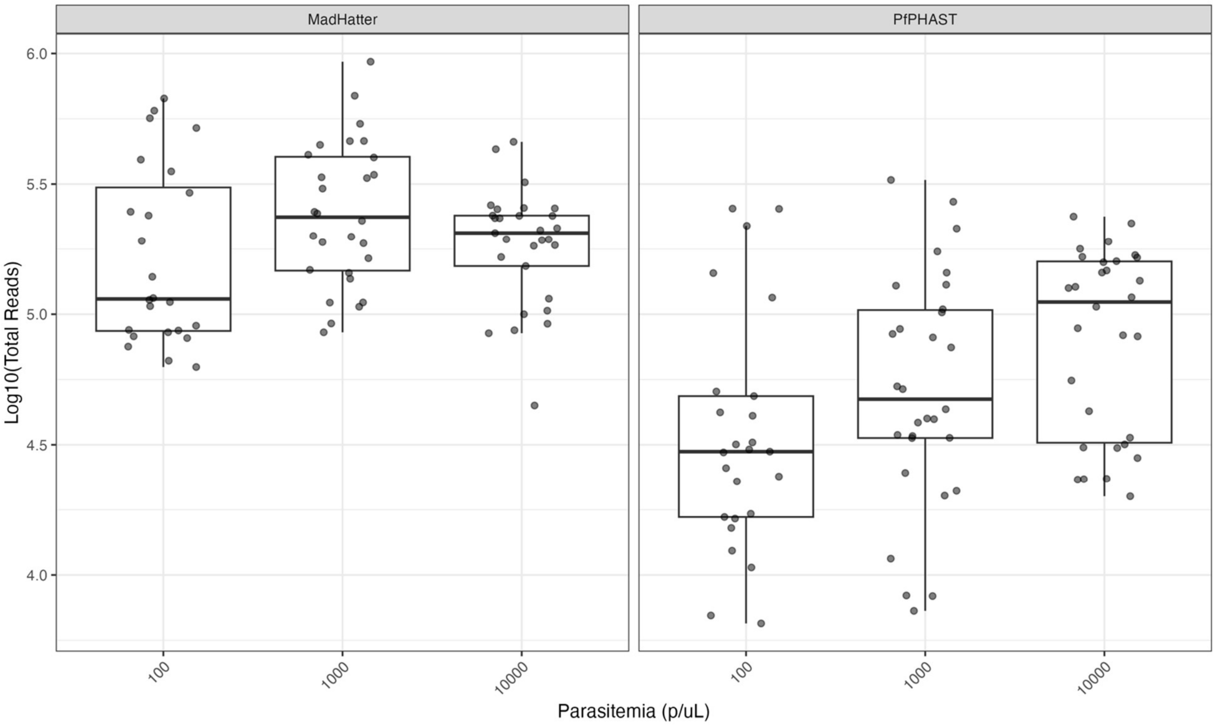


**Supplementary Figure 1. Total sequencing reads per sample across parasite densities for MAD$^4$HatTeR and PfPHAST.** Boxes show the median and interquartile range across samples, with whiskers extending to 1.5× the interquartile range; points are individual samples. Reads were pooled across pairs of replicate MAD$^4$HatTeR libraries, as described in the **Methods**. Parasitemia is expressed in parasite/µL (p/uL).

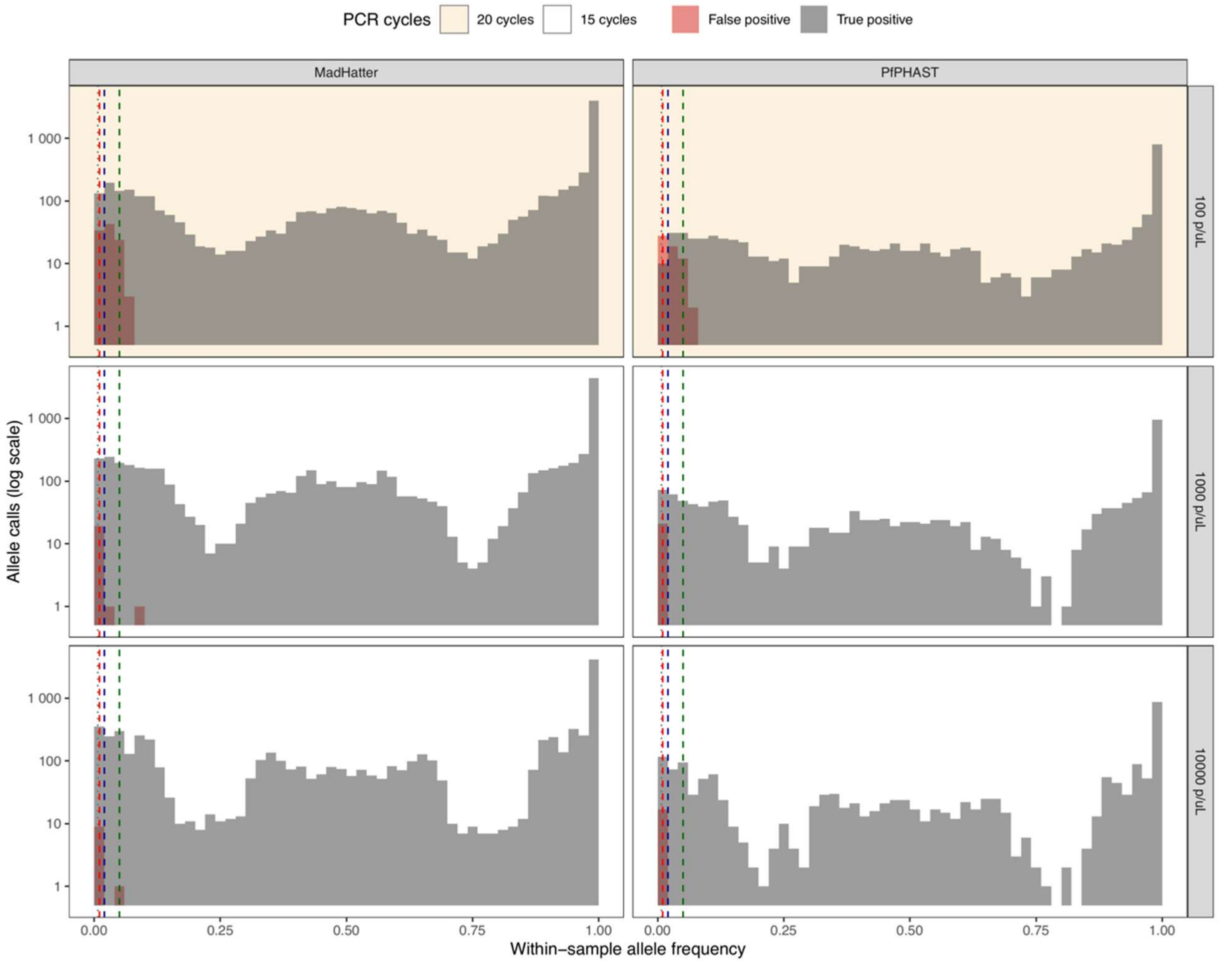


**Supplementary Figure 2. Distribution of true- and false-positive microhaplotype allele calls by within-sample allele frequency (WSAF).** Distribution of true- and false-positive microhaplotype allele calls by within-sample allele frequency (WSAF). Data span the microhaplotype and CSP targets of each panel, which differ between panels, across all control types (two-strain dilution series, the five-strain control, and the D10 and HB3 single-strain controls). Calls required at least 10 reads and 0.75% WSAF. Vertical lines mark the WSAF threshold applied in this analysis (dotted, 0.75%) and alternative thresholds at 1%, 2% and 5% (dashed). Parasitemia is expressed in parasite/µL (p/uL).

**Supplementary Note 1. Per-target design notes and marker exclusions**

***MAD⁴HatTeR resistance module revisions***

The MAD$^4$HatTeR resistance module was revised from pool R1.1 to R1.2 to reduce primer dimerisation, as described in the original publication,[1] by dropping targets not required for routine surveillance: five targets in the chromosome 11 region commonly duplicated in *pfhrp3*-deleted parasites; *pfripr* (codons 511, 673, 755, 1039); *pfcyrpa* (339); *pfrh5* (147, 221, 269, 350, 354, 357, 362); *pfp113* (106, 107, 234); *pfaat1* (135, 162, 185, 230, 238); *pfcrt* (271, 326); *pfpph* (1157); plasmepsin I and plasmepsin IV; and *pfmdr2* copy number targets. One *pfpank1* target was retained as a long (>275 bp) amplicon serving as an internal control. These exclusions carried over into PfPHAST.

***Marker classification***

The WHO compendium of antimalarial drug resistance markers was not available at the time of panel design. Marker selection was therefore based on the evidence available then, and *pfkelch13* targets covered all positions classified as validated or candidate markers of artemisinin partial resistance at that time. Markers added in the compendium published subsequently will be included in the next PfPHAST iteration (primer pools M1.2 and M2.2).

***Targets added in PfPHAST***

Two targets downstream of *pfhrp2* (Pf3D7_08_v3-1378023-1378202 and Pf3D7_08_v3-1382263-1382472) were added to improve deletion breakpoint classification.

One *pfmdr1* target (Pf3D7_05_v3-0959211-0959413) was added. It covers codon 500, where PfMDR1 Y500N has been proposed as a candidate marker of lumefantrine tolerance following the observation of decreased parasite susceptibility to both dihydroartemisinin and lumefantrine in northern Uganda.[2] Evidence for this marker remains limited and it does not meet the criteria applied to the other resistance markers included here, but the target was retained because the additional amplicon also improves *pfmdr1* copy number estimation.

***Targets redesigned***

Five *pfkelch13* targets were redesigned with new primers to extend coverage to previously missed amino acid positions.

| **PfPHAST target** | **Pool** | **Replaces MAD$^4$HatTeR target** |
|---|---|---|
| Pf3D7_13_v3-1724844-1725047 | M1.1 | Pf3D7_13_v3-1724925-1725137 |
| Pf3D7_13_v3-1725038-1725224 | M2.1 | Pf3D7_13_v3-1725151-1725336 |
| Pf3D7_13_v3-1725216-1725423 | M1.1 | Pf3D7_13_v3-1725231-1725441 |
| Pf3D7_13_v3-1725417-1725633 | M2.1 | Pf3D7_13_v3-1725403-1725620 |
| Pf3D7_13_v3-1725489-1725698 | M1.1 | Pf3D7_13_v3-1725493-1725694 |

***Targets reassigned between pools***

MAD$^4$HatTeR includes overlapping amplicons for the *pfcrt* and *pfhrp2* regions in both reactions. In each case the reaction 1 target consistently underperformed, so the reaction 2 target was retained and moved to pool M1.1: for *pfcrt*, Pf3D7_07_v3-0403507-0403717 was retained in place of Pf3D7_07_v3-0403499-0403683; for *pfhrp2*, Pf3D7_08_v3-1375237-1375418 in place of Pf3D7_08_v3-1375056-1375259.

***Targets and markers excluded***

*pfcsp*: Pf3D7_03_v3-0221217-0221388 was excluded, as it covers no sequence variation in global whole genome sequencing data and its coding sequence partially overlaps a retained target. The two retained targets cover the C-terminal domain, including the Th2R, Th3R and DV10 epitopes.

*pfcrt* and *pfmdr1*: codons considered not relevant to African parasite populations at the time of design were not included. These are *pfcrt* 218, 220, 343, 350 and 353, and *pfmdr1* 371 (Pf3D7_05_v3-0958952-0959135) and 1034 and 1042 (Pf3D7_05_v3-0960841-0961021). All will be included in the next PfPHAST iteration (primer pools M1.2 and M2.2).

*pfdhfr*: Pf3D7_04_v3-0748617-0748789 was redundant, as all marker codons in that region are covered by a retained amplicon.

*pfkelch13*: N-terminal domain targets were not included. All positions classified as validated or candidate markers at the date of design are covered by the retained targets.

Plasmepsins: Pf3D7_14_v3-0294534-0294741 (plasmepsin II) and Pf3D7_14_v3-0297960-0298129 (plasmepsin III) were not required for copy number estimation.

Chromosome 11: four targets in the region duplicated in parasites carrying the *pfhrp3* deletion (Pf3D7_11_v3-1902070-1902253, Pf3D7_11_v3-1950459-1950650, Pf3D7_11_v3-1968082-1968268, Pf3D7_11_v3-2001757-2001969) are present in MAD$^{4}$HatTeR and were not retained.

*ldh*: the *P. falciparum* target was not retained, as *P. falciparum* presence is established by the rest of the panel.

***Target composition by gene***

*pfcsp* (2 targets), *pfdhfr* (2), *pfmdr1* (4), *pfcrt* (1), *pfdhps* (3), *pfcoronin* (2), *pfkelch13* (5), *pfhrp2* (4: one within the gene and three downstream), *pfhrp3* (5: one within the gene, three upstream and one downstream), plasmepsin II and plasmepsin III (4), *ldh* (4). Codons covered are listed in **Table 1** and full coordinates in **Supplementary Table 1**.

***Microhaplotype loci***

The 20 unlinked loci with the highest minimum expected heterozygosity were selected from MAD$^4$HatTeR pool D1.1 and divided evenly between pools M1.1 and M2.1 (**Supplementary Table 2**).

***Optional add-on pool***

Pool M1.addon contains four mitochondrial *cytb* targets, one each for *P. vivax*, *P. ovale*, *P. malariae* and *P. knowlesi*, and is compatible with both M1.1 and M2.1. *P. ovale curtisi* and *P. ovale wallikeri* are recognised as distinct, non-recombining species; the *P. ovale* target amplifies both. The *P. malariae* target is predicted also to amplify *P. ovale wallikeri*, so a *P. malariae* call from this pool requires confirmation where *P. ovale* may be present.

***Targets excluded from the present analysis***

Six of the 56 PfPHAST targets were excluded from the analyses reported here: the four *ldh* species identification targets, uninformative in *P. falciparum* controls, and two *pfhrp3* targets exceeding 275 bp (Pf3D7_13_v3-2841410-2841661 and Pf3D7_13_v3-2844354-2844599). Fourteen of the 243 MAD$^4$HatTeR targets were excluded on the same criteria, leaving 50 and 229 targets respectively.

***References***


1. Aranda-Díaz, A. *et al.* Sensitive and modular amplicon sequencing of *Plasmodium falciparum* diversity and resistance for research and public health. *Sci. Rep.* **15**, 10737 (2025).
2. Tumwebaze, P. K. *et al.* Decreased susceptibility of *Plasmodium falciparum* to both dihydroartemisinin and lumefantrine in northern Uganda. *Nat. Commun.* **13**, 6353 (2022).
