## Supplementary File 2 for "PfPHAST: *Plasmodium falciparum* Public Health Amplicon Sequencing Tool, a Streamlined Panel for Malaria Genomic Surveillance"

### Multiplexed Amplicons for Drugs, Diagnostics, Diversity and Differentiation using High Throughput Targeted Resequencing (MAD$^4$HatTeR) and *P. falciparum* Public Health Amplicon Sequencing Tool (PfPHAST)

#### Didactic Standard Operating Procedure

Adapted from CleanPlex amplicon sequencing technology by Paragon Genomics.
Version 5 - April 15th 2026
Contributors: Bienvenue Nsengimaana, Innocent Wiringilimaana, Thomas Katairo, Milya Davlieva, Andrés Aranda-Díaz

The multiplexed PCR NGS panel and protocol described in this document was developed at the EPPIcenter at the University of California San Francisco, and it is an adaptation of the CleanPlex amplicon sequencing technology by Paragon Genomics. The publication can be found here.
Please cite as:
Aranda-Díaz, A., Neubauer Vickers, E., Murie, K. et al. Sensitive and modular amplicon sequencing of Plasmodium falciparum diversity and resistance for research and public health. Sci Rep 15, 10737 (2025). https://doi.org/10.1038/s41598-025-94716-5

More information on Paragon Genomics's product can be found at:
www.paragongenomics.com/targeted-sequencing/amplicon-sequencing/cleanplex-ngs-amplicon-sequencing/

The current protocol for MAD$^4$HatTeR/PfPHAST and other relevant information can be found at:
eppicenter.ucsf.edu/resources

| Category | MAD4HatTeR | PfPHAST |
|---|---|---|
| Diverse loci | 165 (microhaplotypes) | 20 (microhaplotypes) |
| ldh or cytb gene in *Plasmodium* spp. | 5 (5 species, incl. *Pf*) | 4 (4 species) |
| Drug resistance (SNPs and CNV) | 38 (15 genes) | 21 (9 genes) |
| Diagnostic resistance (*Pfhrp2/3*) | 30 (3 regions, incl. ch11) | 9 (2 regions) |
| Others (e.g. *Pfcsp*) | 5 (3 genes) | 2 (1 gene) |
| Total targets: | 243 | 56 |

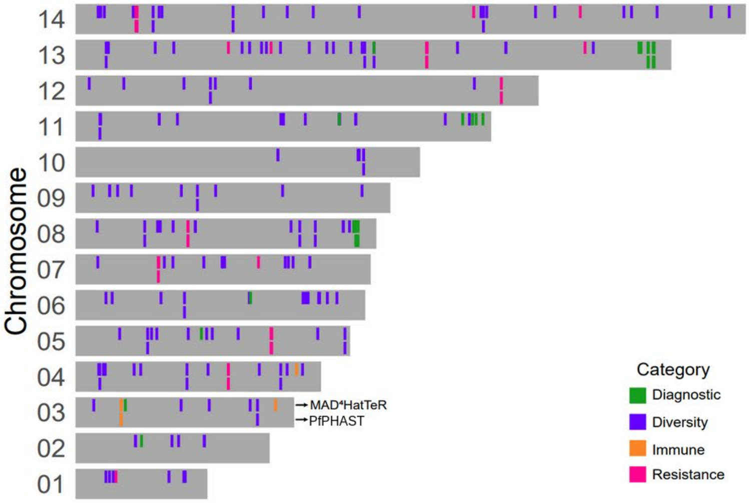

Figure: Summary and genomic location of target loci (using D1.1 + R1.2 + R2.1 configuration for MAD$^4$HatTeR and M1.1 + M2.1 for PfPHAST).

#### Primer pools:

##### MAD⁴HatTeR

MAD[4]HatTeR is a modular assay. Samples can be processed with either or both of two modules: Diversity or Resistance$^+$.

**Diversity:** The diversity module consists of loci of high heterozygosity in available whole genome data. The diversity module can be amplified with primer pool D1.1 (previously 1A)*

**Resistance$^+$:** The resistance$^+$ module consists of loci in drug resistance markers, the *hrp2*/*3* genes associated with diagnostic resistance, and other loci of interest for immune response studies (e.g. *csp*).

The full resistance$^+$ module should be amplified with 2 primer pools (R1.1 and R2.1) that are complementary and incompatible in a multiplexed PCR reaction due to loci overlapping. Pool R1.2 is a subset of R1.1 that significantly reduces primer dimer formation and is also incompatible with pool R2.1. Pools R1.1 and R1.2 contain most of the molecular markers of interest, and thus should take precedence over pool R2.1 if only one mPCR reaction is desired.

The EPPIcenter currently recommends using primer pools R1.2 and R2.1 for drug and diagnostic resistance, and *csp*.

##### PfPHAST

PfPHAST contains a subset of the primers in MAD[4]HatTeR (with some modifications). It is also a modular assay. Primers are distributed in 2 pools (M1.1 and M2.1). These pools contain both diversity and resistance targets but only include validated or prioritary resistance markers, csp C-terminal domain and 20 total microhaplotypes included primarily to correct therapeutic efficacy study outcomes.

A summary of the currently tested primer pools can be found here:

| Module | Pool | Intended reaction | Version | Previous name | Incompatible with (overlaps) | Subset of | Number of targets | Contents |
|---|---|---|---|---|---|---|---|---|
| **Diversity** | **D1.1** | 1 | 1 | 1A | | | 170 | Targets with high heterozygosity and Plasmodium spp. identification |
| **Resistance$^+$** | **R1.1** | 1 | 1 | 1B | R2.1, R1.2 | | 82 | Drug and diagnostic resistance, Plasmodium spp. identification, immune-related targets |
| **Resistance$^+$** | **R2.1** | 2 | 1 | 2 | R1.1, R1.2 | | 31 | Drug and diagnostic resistance, Plasmodium spp. identification, immune-related targets. Complements R1.1 for missing codons |
| **Resistance$^+$** | **R1.2** | 1 | 2 | 5 | R2.1, R1.1 | R1.1 | 47 | Drug and diagnostic resistance, Plasmodium spp. identification, immune-related targets. Subset of R1.1 to reduce primer dimer and increase sensitivity |
| **PfPHAST** | **M1.1** | 1 | 1 | - | M2.1, R1.1, R1.2, R2.1 | | 41 | Drug and diagnostic resistance (validated or prioritary), Plasmodium spp. Identification, csp, and 10 highly diverse microhaplotypes. |
| **PfPHAST** | **M2.1** | 2 | 1 | - | M1.1, R1.1, R1.2, R2.1 | | 15 | Drug and diagnostic resistance (validated or prioritary), Plasmodium spp. Identification, csp, and 10 highly diverse microhaplotypes. |

| | | | | | | | | |
|---|---|---|---|---|---|---|---|---|
| **PfPHAST** | **M1.addon** | 1 or 2 | 1 | - | | | 4 | *cyt-b* targets in non-falciparum *Plasmodium* spp. For more sensitive identification. |

A full list with primer sequences, genomic locations, and genes can be found in [eppicenter.ucsf.edu/resources](eppicenter.ucsf.edu/resources) in the “Pool details” link in the MAD$^4$HaTTeR section.

**Naming convention**

As of **January 2025**, pools have been renamed to allow for consistency and future versions. These changes were also incorporated to release v1.0.0 of the bioinformatic pipeline.
Primer pools names follow a standardized nomenclature: [Module][Reaction].[Version]
Module (letter): indicates the primary use case — D (Diversity), R (Resistance+), or M (PfPHAST).
Reaction (first number): indicates the intended reaction, distinguishing pools that cannot be combined due to overlapping amplicons (e.g., R1.x vs. R2.x).
For example, R1.2 refers to the Resistance+ module, intended for reaction 1, version 2. The addon suffix (e.g., M1.addon) denotes a supplementary pool designed to be spiked into either reaction. A crosswalk between old and new names is provided in the table above.

Diversity pools 3 and 4 were subsets of D1.1. Pool 3: 153 targets with high heterozygosity and Plasmodium spp. identification. Pool 4: 50 targets with high heterozygosity and Plasmodium spp. Identification, a subset of D1.1 to increase throughput. These pools have never been used and thus were not assigned names under the new convention.

#### Laboratory Layout

**Objective**
To minimize contamination of reagents and reactions

We suggest having 3 dedicated spaces for library preparation:

1. ***DNA-free Laminar Flow Hood in Pre-PCR room***
2. ***Laminar Flow Hood in Pre-PCR room***: here we add DNA. Some labs do this in a bench in a Pre-PCR room, some do it on an amplicon-free Laminar Flow Hood within a Post-PCR room.
3. ***Bench in PCR room***

Examples of laboratory layouts:

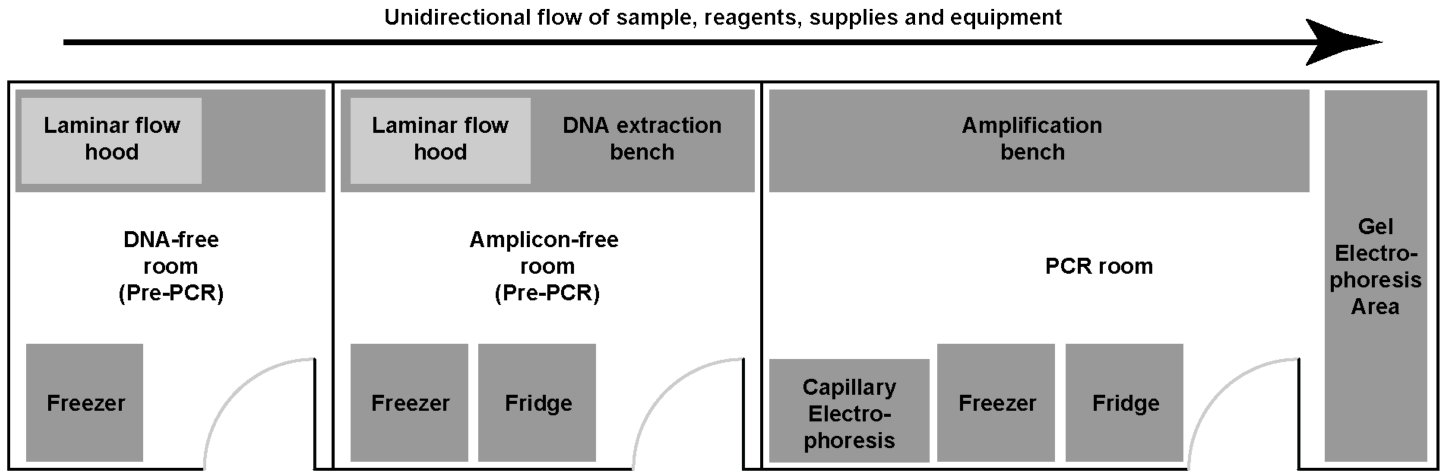

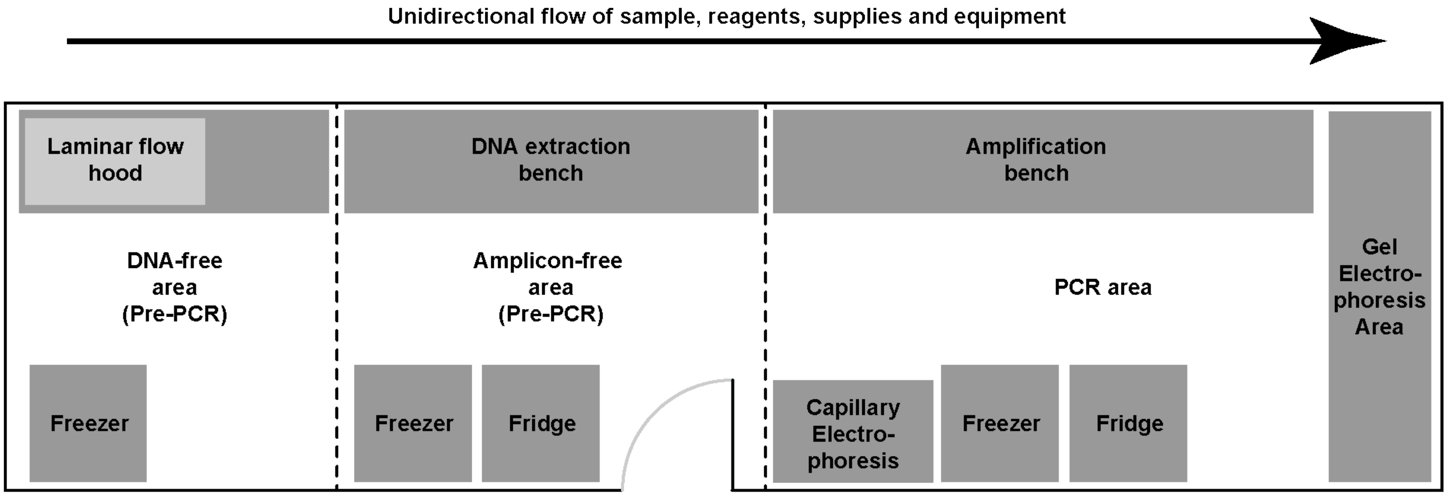

Physical separation between areas (having dedicated rooms) is optimal. The most important feature is the unidirectional flow: no materials go back from PCR to Pre-PCR and from DNA extraction to DNA-free. This is meant to prevent contaminations (specially from highly concentrated amplicons) in reagents and samples that will lead to spurious results that are hard to detect. These layouts imply that **each area will have dedicated equipment** (pipettes, freezers, centrifuges, spinners, etc).

#### Protocol overview

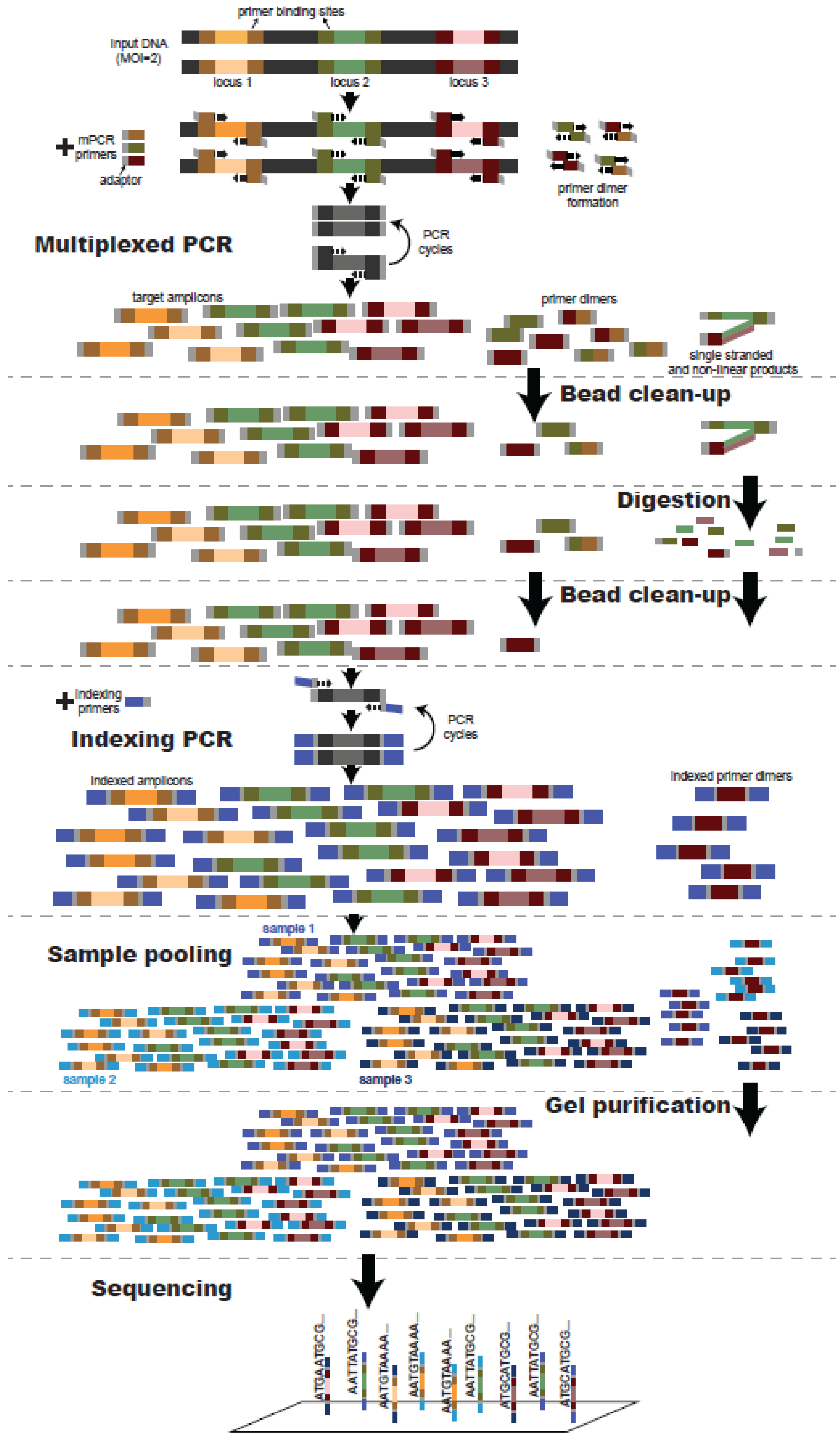

#### PCR programs

Save the following programs in a thermal cycler before starting

**mPCR (multiplex PCR)**

- Initial denaturation: 95 °C 00:10:00
- Amplification for X number of cycles **(refer to table below):**
  - Denaturation: 98 °C 00:00:15 with ramping 3 °C/s
  - Annealing/Extension: 60 °C 00:05:00 with ramping 2 °C/s
- Hold at 10 °C
- Volume: 10 µL
- Lid: 105 °C

| **Parasitemia** | **Total number of mPCR cycles** |
|---|---|
| ≥100 p/µL | 15 |
| <100 p/µL | 20 |

We recommend saving 2 programs with different names (mPCR15 and mPCR20) and protecting them with password or read-only settings

**Digestion**

- 37 °C infinite hold*
- 37 °C for 00:10:00
- Volume: 20 µL
- Lid: OFF

* The infinite hold is used to pre-heat the block to 37 °C to have it ready to start the digestion when the samples are mixed with the digestion reagent. In some thermal cyclers, the infinite hold can be exited with a “next step” function. You will need to figure out how your instrument works.

We recommend saving the program and protecting it with password or read-only settings

**iPCR (indexing PCR)**

- Initial denaturation: 95 °C 00:10:00
- Amplification for 15 cycles:
  - Denaturation: 98 °C 00:00:15 with ramping 3 °C/s
  - Annealing/Extension: 60 °C 00:01:15 with ramping 2 °C/s
- Hold at 10 °C
- Volume: 40 µL
- Lid: 105 °C

We recommend saving the program and protecting it with password or read-only settings

#### Personal protective equipment

PPE should be used in accordance with your laboratory regulations. Proper use of PPE also minimizes sample contamination. Special attention should be paid to powder free glove usage and we recommend changing them frequently

#### Multiplexed PCR (mPCR)

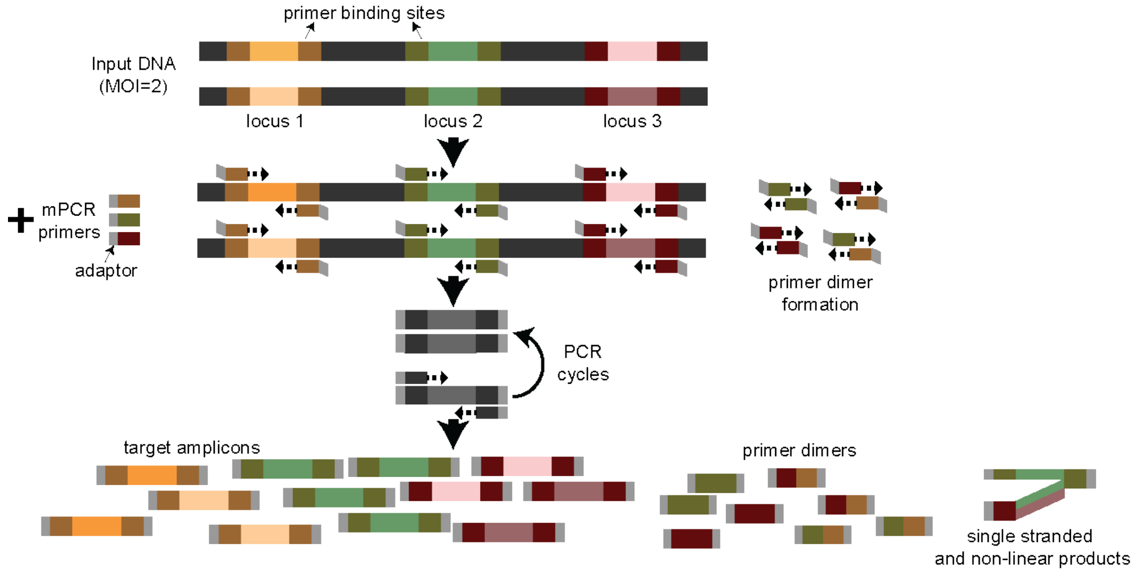

**Objective**

Amplify target regions from *P. falciparum* genomes in 1 or 2 reactions. 2 reactions are needed if using incompatible primer pools, and they can be combined once the targets have been amplified

***In Laminar Flow Hood in Pre-PCR room***

- Thaw primer pools at room temperature. Keep on ice after thawing
- If expecting to run full protocol in one day, thaw CP Reagent Buffer (white tube) and keep on ice after thawing
- Bring 5X mPCR Master Mix (green tube) into PCR Workstation in a cold rack
- Combine the following volumes to prepare the **mPCR reaction mix**. Keep mix on ice or cold rack. Vortex reagents to mix and briefly spin down before opening.

**Generic recipe (add 10% for dead volumes):**
The total volume per reaction is 10 µL, including the mPCR reaction mix and the

| Reagent | Volume (≥100 p/µL) | Volume (<100 p/µL) | … x # reactions x 1.10 (10% extra*) |
|---|---|---|---|
| 5X mPCR Master Mix (green tube) | 2 µL | 2 µL | |
| Each primer pool | 0.5 µL | 0.25 µL | |
| Nuclease-free $H_2O$<br>If you are using less than 6 µL of input DNA, increase water volume (e.g. if using 3 µL DNA, add water up to 7 µL) | Up to 4 µL | Up to 4 µL | |

*: Depending on your setup and expected dead volumes you may want to factor up to 15% extra

**MAD$^4$HatTeR:**

**Note: pools R1.1/R1.2 and R2.1 are incompatible.**

**The following recipes are for 2 reactions/sample, one with pools D1.1+R1.2 and one with R2.1.**

Select the range of Parasitemia for your samples on the plate batch

≥ 100 p/uL (15 mPCR cycles and the normal primer concentration)

< 100 p/uL (20 mPCR cycles and half the normal primer concentration)

**For primer pools D1.1+R1.2**

| **Reagent** | **Volume (≥100 p/µL)** | **Volume (<100 p/µL)** | **… x # reactions x 1.10 (10% extra*)** |
|---|---|---|---|
| 5X mPCR Master Mix (green tube) | 2 µL | 2 µL | |
| Primer pool D1.1 | 0.5 µL | 0.25 µL | |
| Primer pool R1.2 | 0.5 µL | 0.25 µL | |
| Nuclease-free $H_2O$ | 1 µL (or 4 µL if using 3 µL input DNA) | 1.5 µL (or 4.5 µL if using 3 µL input DNA) | |

*: Depending on your setup and expected dead volumes you may want to factor up to 15% extra

**For primer pool R2.1**

| **Reagent** | **Volume (≥100 p/µL)** | **Volume (<100 p/µL)** | **… x # reactions x 1.10 (10% extra*)** |
|---|---|---|---|
| 5X mPCR Master Mix (green tube) | 2 µL | 2 µL | |
| Primer pool R2.1 | 0.5 µL | 0.25 µL | |
| Nuclease-free $H_2O$ | 1.5 µL (or 4.5 µL if using 3 µL input DNA) | 1.75 µL (or 4.75 µL if using 3 µL input DNA) | |

*: Depending on your setup and expected dead volumes you may want to factor up to 15% extra

**PfPHAST:**

**Note: Pools M1.1 and M2.1 are incompatible.**

**The following recipes are for 2 reactions/sample, one with pool M1.1 and one with M2.1.**

**For primer pool M1.1**

| **Reagent** | **Volume (≥100 p/µL)** | **Volume (<100 p/µL)** | **… x # reactions x 1.10 (10% extra*)** |
|---|---|---|---|
| 5X mPCR Master Mix (green tube) | 2 µL | 2 µL | |
| Primer pool M1.1 | 0.5 µL | 0.25 µL | |
| Nuclease-free $H_2O$ | 1.5 µL (or 4.5 µL if using 3 µL input DNA) | 1.75 µL (or 4.75 µL if using 3 µL input DNA) | |

*: Depending on your setup and expected dead volumes you may want to factor up to 15% extra

**For primer pool M2.1**

| **Reagent** | **Volume (for ≥100 p/µL)** | **Volume (<100 p/µL)** | **… x # reactions x 1.10 (10% extra*)** |
|---|---|---|---|
| 5X mPCR Master Mix (green tube) | 2 µL | 2 µL | |
| Primer pool M2.1 | 0.5 µL | 0.25 µL | |
| Nuclease-free $H_2O$ | 1.5 µL (or 4.5 µL if using 3 µL input DNA) | 1.75 µL (or 4.75 µL if using 3 µL input DNA) | |

*: Depending on your setup and expected dead volumes you may want to factor up to 15% extra

- Vortex mixes, briefly spin and keep on ice
- Aliquot 4 µL (or 7 µL if using 3 µL input DNA) mPCR mix into PCR tubes/wells (single tubes, strips or plate). [1]
  Keep tubes on ice
- Put primers and 5X mPCR Master Mix back in freezer
- Transfer tubes to ***Laminar Flow Hood in Pre-PCR Room***

***In Laminar Flow Hood in Pre-PCR room***

- Add 6 µL (or 3 µL) DNA sample to each labeled tube/well, independent of parasitemia.
  Note: total volume is 10 µL
- Vortex and briefly spin down to collect liquids
- Transfer the tubes to the thermal cycler. Double-check that the settings are correct and run the corresponding pre-set **mPCR** program**:**

  Initial denaturation: 95 °C 00:10:00
  Amplification for X number of cycles (15 for ≥100 p/µL, 20 for <100 p/µL):
    Denaturation: 98 °C 00:00:15 with ramping 3 °C/s
    Annealing/Extension: 60 °C 00:05:00 with ramping 2 °C/s
  - Hold at 10 °C
  - Volume: 10 µL
  - Lid: 105 °C

[1] You may do this 1 tube/well at a time, or transfer the volume into a reservoir or PCR tube strip and use a multichannel

#### Preparation of reaction mixes

**After starting the thermocycler protocol prepare for next steps**

- Bring Clean Mag Magnetic Beads and TE buffer to room temperature
- Bring STOP buffer to room temperature and aliquot into PCR tube strip (~200 µL per tube) so that you can use a multichannel
- Make 80% ethanol with nuclease-free water (you will need 800 µL per sample, or 400 µL if stopping after first bead clean-up)

**If you are not stopping at the safe stopping point after the first bead clean-up, also do this:**

- Bring index primers out of the freezer and thaw on ice
- Make a plan for sample indexing. Write down what index you will use for each sample in the table at the end of this worksheet
- Make digestion and indexing PCR mixes

***In DNA-free Laminar Flow Hood in Pre-PCR room***

If you are splitting the protocol in 2 days, make these 2 mixes at the beginning of the second day

- Combine the following volumes to prepare the **Digestion reaction mix:**
  Vortex reagents to mix and briefly spin down before opening.

| **Reagent**[2] | **Volume** | **… x # reactions x 1.10 (10% extra*)** |
|---|---|---|
| Nuclease free water | 6 µL | |
| CP reagent buffer | 2 µL | |
| CP digestion reagent | 2 µL | |

*: Depending on your setup and expected dead volumes you may want to factor up to 15% extra

**Note: Make sure that CP Reagent Buffer is completely dissolved before using**

- Combine the following volumes to prepare the **indexing PCR reaction mix:**
  Vortex reagents to mix and briefly spin down before opening.

| **Reagent**[3] | **Volume** | **… x # reactions x 1.10 (10% extra*)** |
|---|---|---|
| Nuclease free water | 18 µL | |
| 5X 2$^{nd}$ PCR Master Mix | 8 µL | |

*: Depending on your setup and expected dead volumes you may want to factor up to 15% extra

- Put CP reagent buffer and digestion reagent, and 2$^{nd}$ PCR Master Mix back in freezer

#### Multiplexed PCR (mPCR), continued

[2] To use less tips, and if you do not stop at the first safe stopping point, you may add 10 µL TE buffer per reaction
[3] To use less tips you may add 10 µL TE buffer per reaction

***In bench in Post-PCR room***

- Once mPCR reaction is done, remove tubes from the thermal cycler
- **Immediately** add STOP buffer and mix reactions if needed:

**For 2 reactions** (e.g. D1.1+R1.2 and R2.1, or M1.1 and M2.1):
Mix the 2 reactions for each sample. **Be careful not to mix different samples together**

- Add 4 µL STOP buffer to one of the reactions for each sample
- Change the volume in the pipet to 14 µL (or 10 µL if that's the maximum)[4] and transfer all the volume to the corresponding sample tubes for the other reaction so that both reactions and the STOP buffer are combined in one tube
  Note: total volume is 24 µL

**For 1 reaction** (e.g. D1.1.+R1.1 only, or R2.1 only, or M1.1 only, or M2.1 only)[5]:

- Add 2 µL STOP buffer to each of the tubes
- Add 10 µL TE Buffer to each of the tubes
  Note: total volume is 22 µL

[4] You may use the same tips to add STOP buffer and move and combine the 2 reactions for the same sample
[5] To use less tips, you may combine TE and STOP buffers and add together

<u>**Post-mPCR Bead Clean-up**</u>

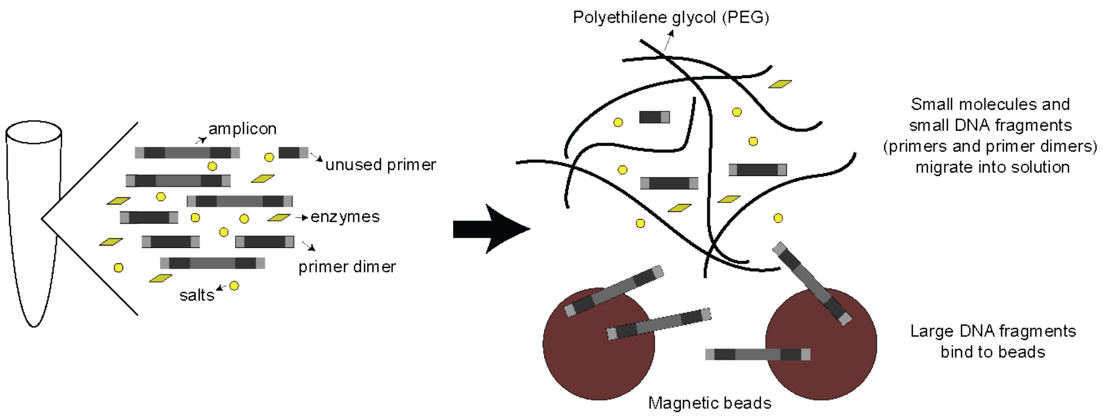

**How are larger DNA molecules captured by the beads?**
Large DNA molecules cannot diffuse into the "mesh"

**Objective**
Remove mPCR reaction leftovers (enzymes, salts and unused primers) and, more importantly, primer dimers from the solution

***In bench in Post-PCR***

- If you haven't, prepare 80% ethanol with nuclease-free water
- Make sure that CleanMag Magnetic Beads are at room temperature and are well mixed
- Add 1.3X bead solution to each tube:
    - 31 µL if you mixed 2 reactions
    - 29 µL if you only had 1 reaction per sample
- Vortex vigorously to mix, or pipet up and down, and spin to collect liquids (not long enough to pellet beads) **After this step, and until resuspension in TE Buffer, do not vortex the mixture and treat it carefully**
- Incubate for 00:05:00 minutes at room temperature
- Place on magnetic stand for 00:03:00 minutes or until the beads are collected on the side of the tubes/wells and the liquid is clear
- Remove all the liquid with a pipet set to >60 µL. Here and in all the following steps, avoid scraping the tube wall where the beads are.
- Briefly spin down*, place back in the magnetic stand and remove the liquid leftovers using a P20 or P10 pipet set to maximum volume. Choose a pipet with thin pipet tips to better take leftovers.
  ***: Place the tubes in the spinner with the beads so that they are on the outside, further away from the center or axis of rotation so that centrifuge force doesn't push them towards the opposite wall**
- Add 180 µL 80 % ethanol
- To wash the beads, rotate the tubes so that the beads migrate from one wall to the other. In magnetic stand for plates with vertical magnets, move the plate or strip by one column.
- Incubate for 00:02:00 or until all beads have migrated. You may need to **CAREFULLY** flick the tubes/wells
- Remove the liquid with a P200 pipet
- Repeat wash: add 180 µL 80% ethanol
- Rotate or move the tubes/plate so that the beads migrate from one wall to the other.
- Incubate for 00:02:00 or until all beads have migrated. You may need to **CAREFULLY** flick the tubes/wells
- Remove the liquid with a P200 pipet
- Briefly spin down, place back in the magnetic stand and remove all the remaining liquid using a P20 or P10 pipet (this is only on the second wash, not the first one)
- Leave tubes/wells open to dry at room temperature. **Generally, a 5 min dry time is enough, but it will depend on room temperature and humidity**. You may not have to wait and just add TE buffer as soon as you finish removing the liquid from the last well

The beads should look matte, not shiny. **Under-drying (carrying ethanol) and over-drying (cracking) can lead to reduced yield**

- Add 10 µL TE buffer equilibrated at room temperature. Close the tubes/wells and vortex vigorously and/or tap to re-suspend the beads.[6]
- Quickly spin down to collect liquids
  The magnetic beads will not affect the rest of the reactions, there is no need to remove them

**Where is my DNA?**
While larger DNA is precipitated out (because of the fine mesh in the polymer or because it is not soluble in ethanol) DNA will be on the

~~~~~~~~~~~~~~~~~~~~~~~~~~~~~~~~~SAFE STOPPING POINT~~~~~~~~~~~~~~~~~~~~~~~~~~~~~~~~~

**If you want to stop here, store at -20 °C**

**If you stopped in the previous step and left samples at -20 °C, at the beginning of day 2:**

- Thaw CP Reagent Buffer (white tube) and keep on ice after thawing
- Bring index primers out of the freezer
- Bring Stop Buffer, TE Buffer and CleanMag Magnetic Beads to room temperature
- Make 80% ethanol with nuclease-free water (400 µL per sample)
- Write down what index you will use for each sample in the table at the end of this worksheet
- Make digestion and indexing PCR mixes in DNA-free Laminar Flow Hood
- Bring samples out of the freezer, thaw and spin down to collect liquids

---

[6] If you are continuing with the full protocol and want to use less tips, you may resuspend directly in a mixture of TE and the digestion reaction mix
~~~~~~~~~~~~~~~~~~~~~~~~~~~~~~~~~

#### Digestion

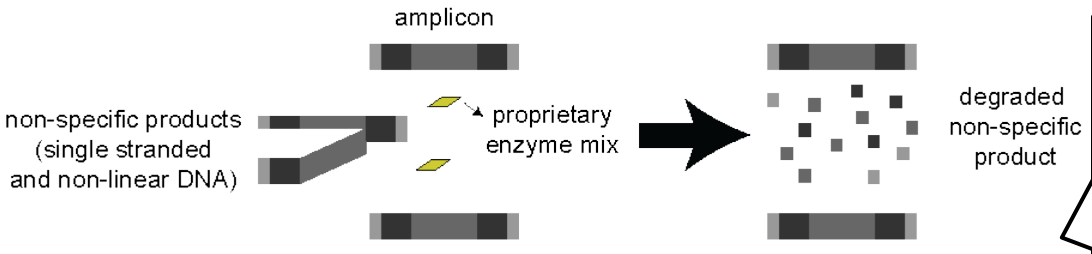

**Why do we need to remove non-specific products?**
Non specific products, such as chimeras of [illegible]

**Objective**
To remove non-specific products that are not double stranded DNA
Warning: if the digestion reaction is not stopped in time, degradation of library products can occur. Promptly add STOP buffer and proceed to bead clean-up.

***In bench in Post-PCR room***

- Start the protocol without the samples. If needed, once the block has reached 37 °C press Pause
- Vortex Digestion reaction mix and briefly spin down to collect liquids
- Add 10 µL of digestion reaction mix to each tube[7]
- Vortex and quickly spin down to collect liquids
- Transfer tubes to thermal cycler pre-set at 37 °C and press Resume (or equivalent option in your thermal cycler) to incubate for 10 min
- After 10 min, **immediately** add 2 µL of Stop Buffer in each tube

[7] To use less tips, and if you did not stop at the previous safe stopping point, you may add the digestion reaction mix together with TE buffer in the last steps of the previous section

#### Post-Digestion Bead Clean-up

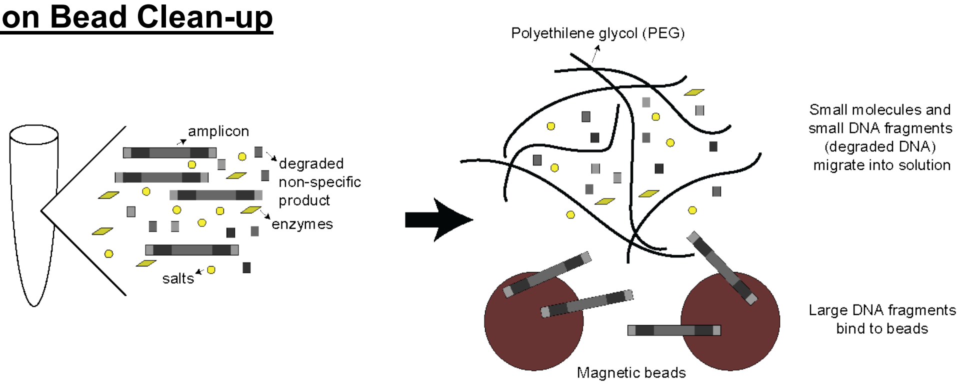

##### Objective

Remove enzymes, salts and short digestion products from the solution

***In bench in Post-PCR room***

- If you haven't, prepare 80% ethanol with nuclease-free water
- Make sure that CleanMag Magnetic Beads are at room temperature
- Add 1.3X bead solution to each tube: 29 µL
- Vortex vigorously, or pipet up and down, to mix and spin to collect liquids (not long enough to pellet beads) **After this step, and until resuspension in TE Buffer, do not vortex the mixture and treat it carefully**
- Incubate for 00:05:00 minutes at room temperature
- Place on magnetic stand for 00:03:00 minutes or until the beads are collected on the side of the tubes/wells and the liquid is clear
- Remove all the liquid: first with a pipet set to >60 µL
- Briefly spin down*, place back in the magnetic stand and remove the liquid leftovers using a P20 or P10 pipet set to maximum volume. Choose a pipet with thin pipet tips to better take leftovers. ***: Place the tubes in the spinner with the beads so that they are on the outside, further away from the center or axis of rotation so that centrifuge force doesn't push them towards the opposite wall**
- Add 180 µL 80 % ethanol
- To wash the beads, rotate the tubes so that the beads migrate from one wall to the other. In magnetic stand for plates with vertical magnets, move the plate or strip by one column.
- Incubate for 00:02:00 or until all beads have migrated. You may need to **CAREFULLY** flick the tubes/wells
- Remove the liquid with a P200 pipet
- Repeat wash: add 180 µL 80% ethanol
- Rotate or move the tubes/plate so that the beads migrate from one wall to the other.
- Incubate for 00:02:00 or until all beads have migrated. You may need to **CAREFULLY** flick the tubes/wells
- Remove the liquid with a P200 pipet
- Briefly spin down, place back in the magnetic stand and remove all the remaining liquid using a P20 or P10 pipet (this is only on the second wash, not the first one)
- Leave tubes/wells open to dry at room temperature. **Generally, a 5 min dry time is enough, but it will depend on room temperature and humidity**. You may not have to wait and just add TE buffer as soon as you finish removing the liquid from the last well. The beads should look matte, not shiny. **Under-drying (carrying ethanol) and over-drying (cracking) can lead to reduced yield**
- Add 10 µL TE buffer equilibrated at room temperature. Close the tubes/wells and vortex vigorously and/or tap to re-suspend the beads. [8]
- Quickly spin down to collect liquids

#### Indexing PCR

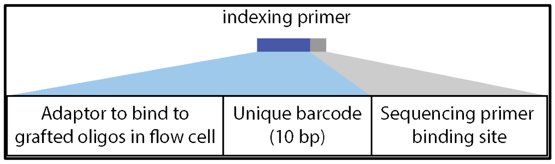

[8] To use less tips, you may resuspend directly into a mixture of TE and indexing PCR mix

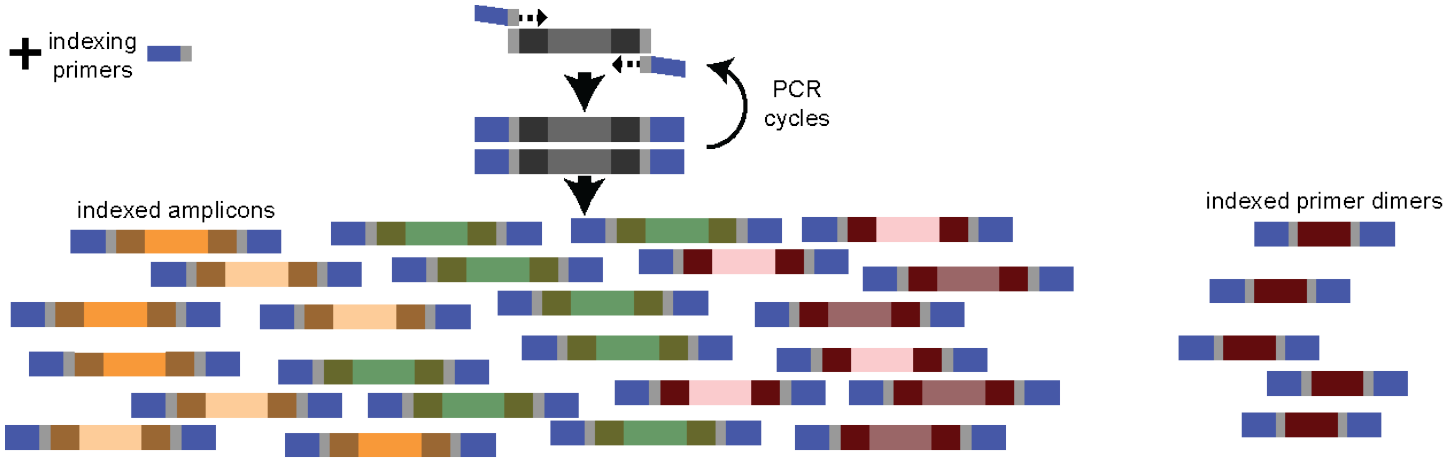

**Objective**
To further amplify amplicons and append adaptors so that the DNA binds to the sequencer flow cell, and a unique DNA barcode in each sample

***In bench in Post-PCR room***

- Vortex Indexing PCR mix and briefly spin down to collect liquids
- Add 26 µL Indexing PCR mix to each tube
- Spin down indexing primer plate and cut out the cover from the wells you will use
- Add 4 µL of indexing primers from a unique index plate well into each of the samples
- Reseal indexing PCR plate
- Vortex and briefly spin to collect liquids
- Double-check that the settings are correct and run the pre-set **iPCR** program**:**

  Initial denaturation: 95 °C 00:10:00
  Amplification for 15 cycles :
    Denaturation: 98 °C 00:00:15 with ramping 3 °C/s
    Annealing/Extension: 60 °C 00:01:15 with ramping 2 °C/s
  - Hold at 10 °C
  - Volume: 40 µL
  - Lid: 105 °C

- Make note of the primers used
- After the program is done, take tubes out and store or keep using

**Why do I need to use different indexes for each sample?**
Indexes are unique DNA sequences (barcodes) that are appended to each molecule. Indexes are sequenced alongside the fragments of DNA

~~~~~~~~~~~~~~~~~~~~~~~~~~~~~~~~~~SAFE STOPPING POINT~~~~~~~~~~~~~~~~~~~~~~~~~~~~~~~~~~

**If you want to stop here, store at -20 °C**
~~~~~~~~~~~~~~~~~~~~~~~~~~~~~~~~~~

#### Capillary electrophoresis

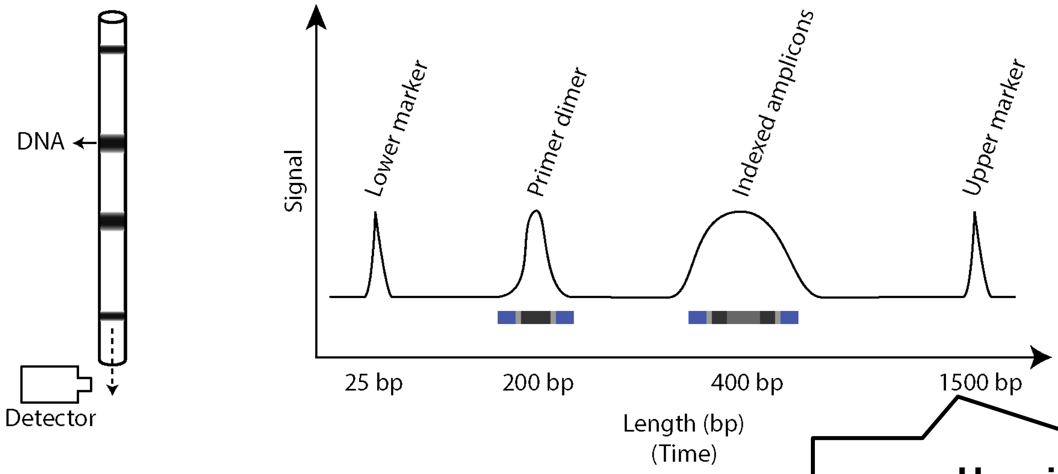

**How is DNA size and concentration determined?**
Just as in regular gel electrophoresis, smaller DNA fragments will travel faster through the polymer when exposed to an electric field. The readouts in [illegible]

**Objective**
To assess the quality of libraries:
1. Do we have indexed amplicons?
2. Do we have primer dimers?
3. What are our library concentrations?
4. Are our negative controls clean (contamination check)

- Briefly spin down tubes and place on magnetic stand to separate beads, which cannot be loaded into capillary electrophoresis systems
- Select random samples to run on capillary electrophoresis. Include negative and positive controls when available
- Follow the instructions for the specific instrument you are using

**Results interpretation:**
1. Internal controls: All samples should have lower and upper markers that come with the buffer. These markers are used to back-calculate the length of DNA fragments based on the time they were detected. If no markers are observed, the readings should not be interpreted
2. Negative control: A negative control with no template for mPCR should still have primer dimers that are amplified and are indexed. A peak should be observed at ~200 bp and no peak should be observed at ~400 bp
3. Positive control (optional) and samples: Positive control and samples should have an amplified library (~400 bp). Many samples will also have a peak at ~200 bp corresponding to the primer dimers that were not eliminated during bead cleanups and were amplified in indexing PCR. Library concentrations would optimally be at ~10 nM.

#### Sample pooling

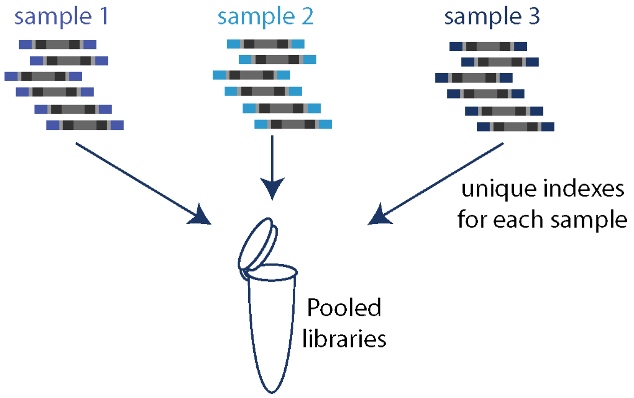

**Objective**
Combine libraries from multiple samples to be sequenced

***In bench in Post-PCR room***

- Before starting make 80% ethanol with nuclease-free water and bring CleanMag Magnetic Beads to room temperature
- Create a sample sheet (you may use the templates available at eppicenter.ucsf.edu/resources)
- Double check that indexes and sample names are not duplicated
- Briefly spin down tubes and place on magnetic stand to separate beads
- Pool samples by mixing them into a single 1.5 mL microcentrifuge tube

  We recommend using the following volumes if using 15 cycles in mPCR:
    - 30 µL for <10 p/µL
    - 20 µL for 10 to <100 p/µL
    - 15 µL for 100 to <1,000 p/µL
    - 6 µL for 1,000 to 10,000 p/µL
    - 3 µL for ≥10,000 p/µL

  If you have samples with unknown parasitemia, but suspect they are high, mix 10 µL of each sample.

  If you have capillary electrophoresis for each of the samples, pool with volumes inversely proportional to the concentration of the 250-650 bp region

  **Total volume in the pool: ……..**

#### Post-Pooling Bead cleanup

**Objective**

Remove leftover primer dimers and indexing PCR reagents

- Add 1X volume of beads to the pool (same volume you entered above). You may need to split in more than 1 tube if the total volume is > 1.5 mL
- Vortex and briefly spin to collect liquids (but don't pellet beads)
- Incubate for 00:05:00 at room temperature
- Place tube(s) in microcentrifuge tube magnetic stand and incubate for 00:03:00 or until the solution is clear
- Remove supernatant
- Briefly spin down and place back in magnetic stand, remove the rest of the supernatant
- Add 1.5 mL 80% ethanol
- Rotate tube(s) and incubate until all beads have migrated to the opposite wall
- Remove ethanol supernatant
- Add 1.5 mL 80% ethanol
- Rotate tube(s) and incubate until all beads have migrated to the opposite wall
- Remove ethanol supernatant
- Briefly spin down, place back in magnetic stand and remove the rest of the supernatant
- Leave tube(s) open to dry at room temperature. **Generally, a 5 min dry time is enough, but it will depend on room temperature and humidity**. The beads should look matte, not shiny. **Under-drying (carrying ethanol) and over-drying (cracking) can lead to reduced yield**
- Resuspend beads with 43 µL TE Buffer.

If using multiple tubes, resuspend in one tube and use that resuspension to resuspend the rest of the tubes so that the final total volume is 43 µL

- Incubate at Room Temperature for 00:02:00
- Place tube in magnetic stand and incubate at room temperature for 00:03:00 or until liquid is clear
- Transfer 40 µL to a clean, labeled tube

**Why is the ratio of beads to DNA lower?**

In previous clean-ups we added 1.3X the volume in beads. This time we use 1X. A lower ratio of bead solution leads to more diluted PEG (the polymer that makes the mesh to exclude large molecules). A lower concentration will allow larger molecules to diffuse into solution as the mesh is less tight.
One of the goals of these cleanups is to remove primer dimers. Before the [illegible]

#### Capillary electrophoresis

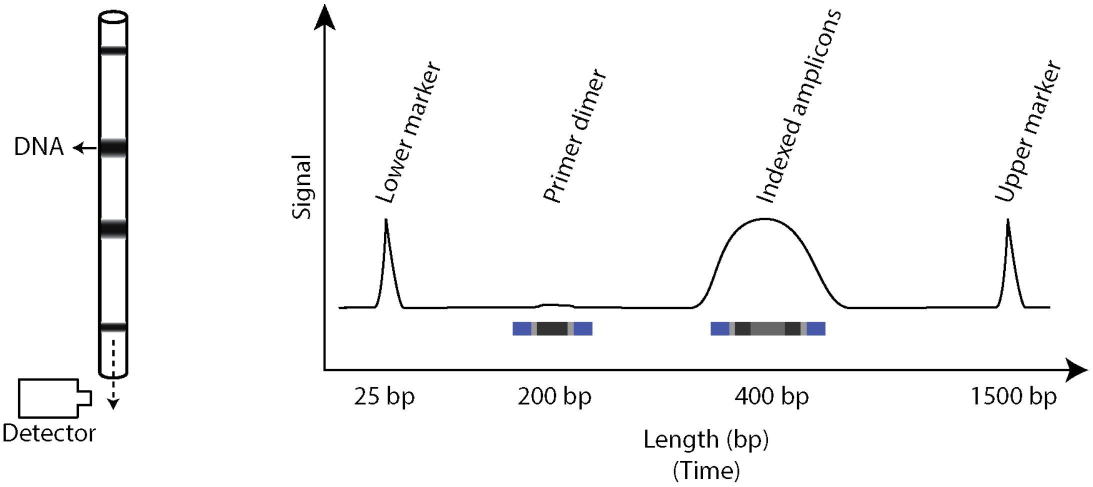

**Objective**
To assess pool quality and determine need of further purification

- Follow the instructions for the specific instrument you are using

**Results interpretation and recommendations:**
The following concentrations, volumes and percentages are points of reference, and could be modified depending on laboratory needs (sequencing requirements, general yield, etc). At the EPPIcenter we have better yields with gel purification than bead cleanup so our preferred method for high dimer contents is gel purification. If the library concentration is high enough, we prefer a bead cleanup for simplicity.

| **Primer dimer** | **400 bp concentration** | **Recommendation**<br>For all, assess quality with capillary electrophoresis and follow these recommendations again |
|---|---|---|
| >5% | >20 nM | Perform a 1X bead clean up and resuspend in 15 µL |
| >5% | <20 nM | Perform gel purification to avoid losing library and more stringently remove primer dimers |
| 1-5% | >50 nM | Perform a 1X bead clean up and resuspend in 15 µL |
| 1-5% | 20-50 nM | Perform gel purification to avoid losing library and more stringently remove primer dimers |
| 1-5% | <20 nM | Proceed with the pool as is, no further purification is recommended to avoid losing the library. You can consider performing a 0.8X bead cleanup and resuspending in 15 µL to concentrate the library. Note that most sequencing cores will require at least 4 nM |
| <1% | <20 nM | Proceed with pool as is, no further purification is recommended to avoid losing library |
| <1% | >20 nM | Perform a 1X bead clean up and resuspend in 15 µL |

#### Gel purification

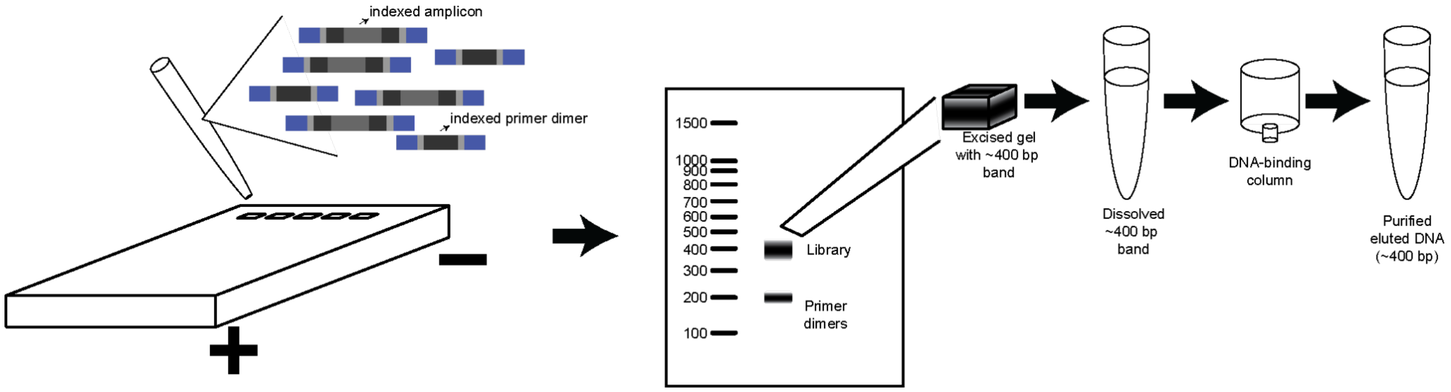

**Objective**

To remove any remaining primer dimers

**Procedure**

1. Cast a 2.5% agarose gel in TAE buffer with 1X SYBRsafe
2. Place the gel in an electrophoresis system and fill up with TAE
3. Load 5-10 µL DNA ladder in one well
4. Add 4 µL 6X loading buffer to 20 µL of the pooled libraries. Vortex and spin to collect liquids
5. Load the pool into 1 or more lanes (depending on comb size you may not be able to fit in one lane). Leave an unused lane between the ladder and the pool
6. Run with constant voltage at 140 V for 1 h
7. After 1 h, quickly image the gel. If the bands (200 and 400 bp) are clearly separated, you may continue to excise. Otherwise, run for longer
8. Excise the 400 bp band
9. Using a DNA gel extraction kit, dissolve the excised gel and run through a column following manufacturer's instructions
10. Elute with 15 µL elution buffer

**Haven't we got rid of primer dimers already?**

In some cases, the previous magnetic bead clean-ups are enough to remove primer dimers completely. But most libraries, specially those with low DNA input or high cycles, carry a lot of primer dimers. A gel extraction is effective as the size selection is more stringent than with PEG, but can only be done with the pooled samples. Otherwise you'd

#### Capillary electrophoresis

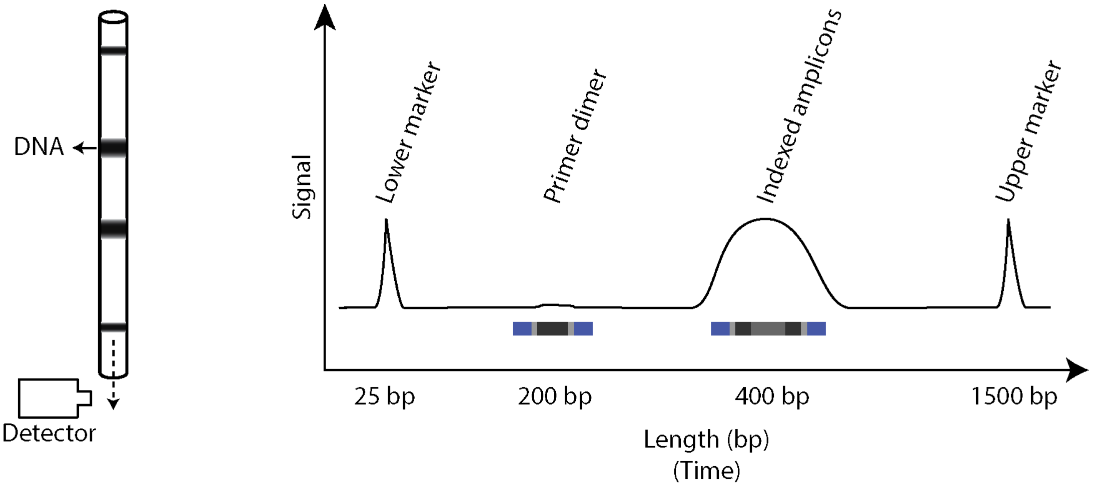

**Objective**
Ensure that primer dimers are at less than 5% of the total sample

- Follow the instructions for the specific instrument you are using

**Results interpretation:**

For the pool to be ready for sequencing, we need to check 2 measurements:

1. The primer dimer peak, if present, should not be more than 5% of the sample. If not, you will need to do another gel purification
2. The libraries should be at ≥ 4 nM (as required per Illumina protocols)

**Was another gel purification necessary? ……..**

**Why do we care so much about primer dimers?**
Primer dimers are fragments of DNA that are barcoded and adapted. Thus, they will attach to the flow cell in the sequencer and get sequenced. But they don't have any information about the sample as they are simply 2 primers that annealed with each other.

#### Sequencing

Now, our libraries consist of DNA fragments that resulted from multiplexed amplification of *P. falciparum* DNA. These libraries have adapters that will allow them to bind to the sequencer's flow cell and start the reactions that lead to sequencing.

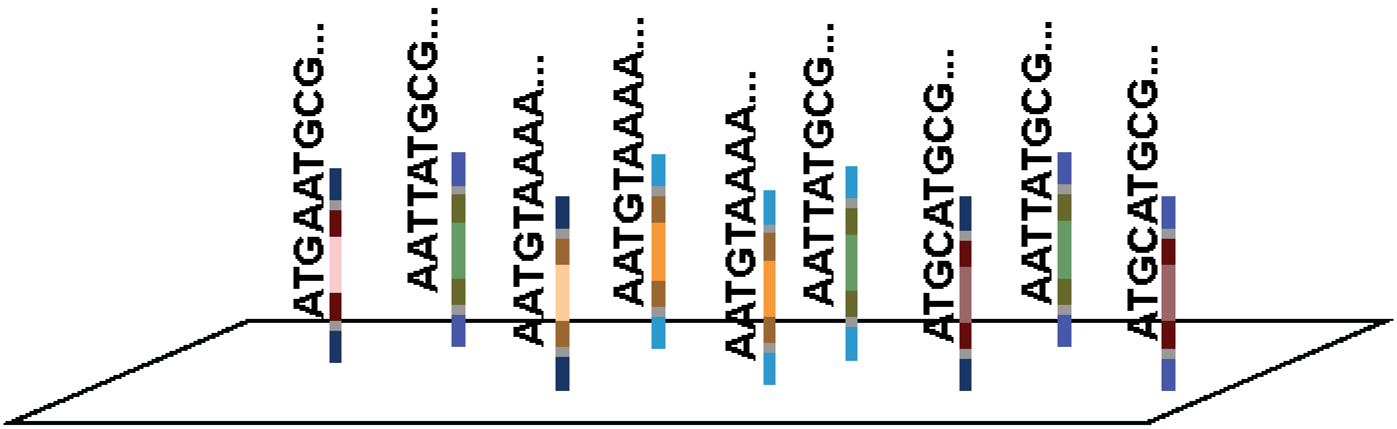

#### Appendix 1: Summary of all sample and indexes used

| Tube No. | Sample id | Primer pools used | mPCR cycle no. | Index used (Set and well) | Volume pooled |
|---|---|---|---|---|---|
